# Mono-ADP-ribosyltransferase 1 (*Artc1*)-deficiency decreases tumorigenesis, increases inflammation, decreases cardiac contractility, and reduces survival

**DOI:** 10.1101/2023.02.06.527366

**Authors:** Hiroko Ishiwata-Endo, Jiro Kato, Hirotake Oda, Junhui Sun, Zu-Xi Yu, Chengyu Liu, Danielle A. Springer, Pradeep Dagur, Martin J. Lizak, Elizabeth Murphy, Joel Moss

**Affiliations:** Pulmonary Branch, National Heart, Lung, and Blood Institute, Maryland, USA; Cardiac Physiology Branch, National Heart, Lung, and Blood Institute, Maryland, USA; Pathology Facility, National Heart, Lung, and Blood Institute, Maryland, USA; Transgenic Core, National Heart, Lung, and Blood Institute, Maryland, USA; Murine Phenotyping Core, National Heart, Lung, and Blood Institute, Maryland, USA; Flow Cytometry Core, National Heart, Lung, and Blood Institute, Maryland, USA; Mouse Imaging Facility, National Institutes of Health, Bethesda, Maryland, USA

## Abstract

Arginine-specific mono-ADP-ribosylation is a reversible post-translational modification; arginine-specific, cholera toxin-like mono-ADP-ribosyltransferases (ARTCs) transfer ADP-ribose from NAD^+^ to arginine, followed by cleavage of ADP-ribose-(arginine)protein bond by ADP-ribosylarginine hydrolase 1 (ARH1), generating unmodified (arginine)protein. ARTC1 has been shown to enhance tumorigenicity as does *Arh1* deficiency. In this study, *Artc1*-KO and *Artc1/Arh1*-double-KO mice showed decreased spontaneous tumorigenesis and increased age-dependent, multi-organ inflammation with upregulation of pro-inflammatory cytokine TNF-*α*. In a xenograft model using tumorigenic *Arh1*-KO mouse embryonic fibroblasts (MEFs), tumorigenicity was decreased in *Artc1*-KO and heterozygous recipient mice, with tumor infiltration by CD8^+^ T cells and macrophages, leading to necroptosis, suggesting that ARTC1 promotes the tumor microenvironment. Furthermore, *Artc1/Arh1*-double-KO MEFs showed decreased tumorigenesis in nude mice, showing that tumor cells as well as tumor microenvironment require ARTC1. By echocardiography and MRI, *Artc1*-KO and heterozygous mice showed male-specific, reduced myocardial contractility. Furthermore, *Artc1*-KO male hearts exhibited enhanced susceptibility to myocardial ischemia-reperfusion-induced injury with increased receptor-interacting protein kinase 3 (RIP3) protein levels compared to WT mice, suggesting that ARTC1 suppresses necroptosis. Overall survival rate of *Artc1*-KO was less than their *Artc1*-WT counterparts, primarily due to enhanced immune response and inflammation. Thus, anti-ARTC1 agents may reduce tumorigenesis but may increase multi-organ inflammation and decrease cardiac contractility.

**Graphical Abstract:** 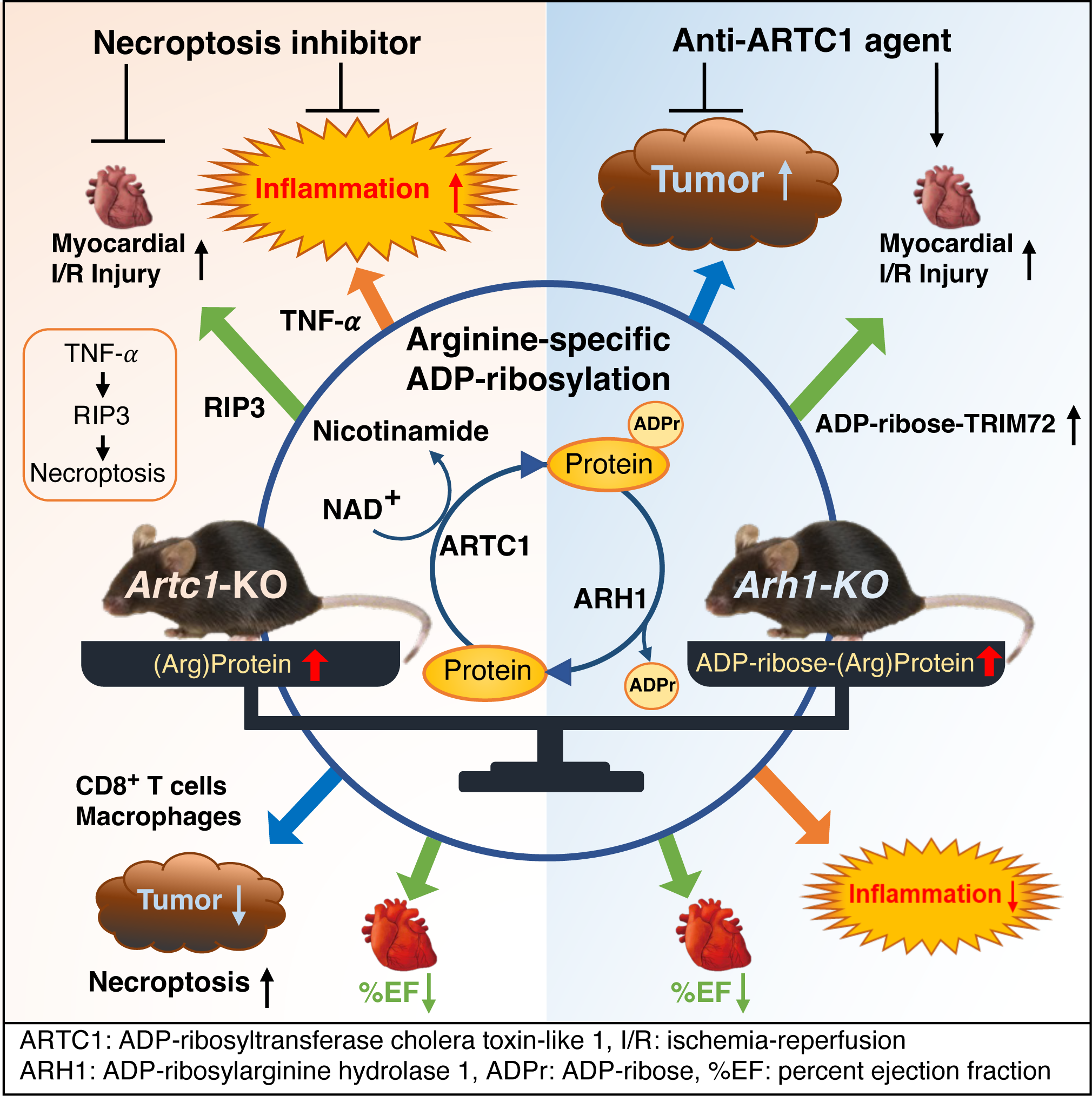

## Introduction

Some bacterial toxins, e.g., cholera, diphtheria, and pertussis toxins, exert their effects on host cells by ADP-ribosylation of critical cellular proteins (1–3). With these bacterial toxins, ADP-ribosyltransferase activity modifies specific amino acids in an acceptor protein, respectively, arginine, diphthamide and cysteine (4–6). In mammalian species, arginine-specific, cholera toxin-like mono-ADP-ribosyltransferases (ARTCs) are expressed on airway epithelium (7), inflammatory cells, e.g., lymphocytes, neutrophils (8), and in various cell types, e.g., muscle and cancer cells (9–13). In sum, mono-ADP-ribosylation is a post-translational modification with functional effects on biological pathways, e.g., innate immunity (14, 15), tumorigenesis (16), oxidative stress (17).

Cholera toxin-like ARTC family and diphtheria toxin-like poly(ADP-ribose)polymerase (PARP) family members possess conserved ART domains, based respectively on R-S-E (18) and H-Y-E (19) sequence motifs. Among mammalian cholera-like ARTC family members, i.e., ARTC1-5, only ARTC1, ARTC2, and ARTC5 catalyze arginine-specific ADP-ribosylation (20). ARTC1 and ARTC2 are attached to the extracellular surface by glycosylphosphatidylinositol (GPI) anchors (10, 21). In human, however, ARTC2 is a pseudogene (18). ARTC5 lacks the signal sequence required for addition of a GPI anchor and is thought to be a secreted exoenzyme (22). NADase activity of ARTC5 is 10 times higher than that of its arginine-specific ADP-ribosyltransferase activity, suggesting that ARTC5 may be acting primarily as an NAD^+^ glycohydrolase or that free arginine is not a good substrate, allowing the NADase activity to predominate (23). Therefore, it appears that ARTC1 is a major contributor to arginine-specific mono-ADP-ribosylation, as has been reported in mouse muscle by comparison of wild-type to *Artc1*-knockout muscle (24).

Mono- and poly-ADP-ribosylations are reversible post-translational modifications of proteins (25–27), DNA (28), and RNA (29). ADP-ribose-acceptor bonds are cleaved by ADP-ribosyl-acceptor hydrolases (ARHs) (25, 30, 31) and macrodomain (MacroD)-containing proteins (32–34). The ARH family comprises three 39-kDa proteins (ARH1-3) that share similar sequence (35). ARH1 and ARH3, however, cleave ADP-ribose-acceptor bonds on different substrates, i.e., ADP-ribosyl-arginine is cleaved primarily by ARH1 (36) and ADP-ribose-serine is cleaved by ARH3 (30, 31). An ARH2 enzymatic activity has not been demonstrated (35, 37).

Macrodomains consist of approximately 190 amino acids and share a somewhat conserved structure; some macrodomains bind mono- and poly-ADP-ribose and cleave ADP-ribose-acceptor bonds (38). The Af1521 macrodomain from archaebacteria *Archaeoglobus fulgidus* binds ADP-ribose and has been used for identification of ADP-ribosylated proteins (24, 39). Macrodomain modules were found in viruses, e.g., SARS-CoV2; bacteria, e.g., *Salmonella typhimurium* (40), *Archaeoglobus fulgidus* (41); and vertebrates, e.g, PARP9, PARP10, PARP14, PARP15 (42–44). In addition to the ADP-ribose binding property of macrodomains, some macrodomains including MacroD1 (45), MacroD2 (45), Af1521 (46–48), DarG (49), PARP10 (42), and TARG1/C6orf130 (50) have been shown to hydrolyze a variety of ADP-ribose-acceptor substrates, e.g., ADP-ribose-glutamate (51), *α*-NAD (47), ADP-ribose-phosphate (38). Thus, Af1521 can both bind ADP-ribose attached to some acceptors and cleave other ADP-ribose-acceptor bonds. The primary substrate recognition for ARH appears to be in the ADP-ribose moiety (47). In agreement, in substrates recognized by ARH1 and ARH3, ADP-ribose attached to both “O” and “N” functional groups is hydrolyzed (37, 52).

A number of diverse proteins are ADP-ribosylated on arginine residues. A recent study found that 354 proteins were ADP-ribosylated on arginine in wild-type mouse heart, mouse skeletal muscle, and C2C12 myotubes (24). Only 6 proteins, however, were ADP-ribosylated on arginine in an ARTC1-independent manner in *Artc1*-KO mouse skeletal muscle (24). Taken together, ARTC1 appears to be the major source of arginine-specific ADP-ribosylation in human and murine skeletal and heart muscle (24). The relative importance in arginine ADP-ribosylation by surface ARTC1 versus that of the secreted ARTC5 may be one of tissue and cell-type specificity. In agreement, proteins ADP-ribosylated by ARTC1, with arginine being the acceptor, have been detected in several mammalian tissues in vivo including human neutrophil peptide-1 (HNP-1) in lung disease, e.g., asthma, idiopathic pulmonary fibrosis (9, 53) and murine tripartite motif-containing protein 72 (TRIM72) in ischemia-reperfusion injury (17).

Other studies have shown that mono-ADP-ribosylation participates in tumor progression, e.g., PARP-7 (54) and PARP-10 (55) promote cell proliferation and tumor progression in cancer cells. A mouse xenograft model showed that increased ARTC1 protein levels were associated with epithelial-mesenchymal transition (EMT), cell proliferation, and angiogenesis, suggesting that ARTC1 may serve as a tumor promotor (56, 57). In contrast, ARH1 appears to be a tumor suppressor (16, 58). In humans, *ARTC1* mRNA and protein levels are increased in human lung adenocarcinoma (13) and glioma tissues (59). These studies have resulted in this proposal that anti-ARTC1 antibodies or other agents might be used as anti-tumor therapeutics (13, 59).

To gain a better understanding of an anti-ARTC1 agent, we used CRISPR/Cas9 to generate an *Artc1*-KO mouse to study *Artc1* deficiency and potential effects of an anti-ARTC1 agent. We found that pathways including ARTC1 and ARH1 are found in multiple organs, and interact in multiple biological events. *Artc1*-KO and *Artc1*-heterozygous (HT) mice exhibited decreased tumor formation with enhanced infiltration of CD8^+^ T cells and macrophages, suggesting that ARTC1 promotes a tumor microenvironment. However, simultaneously in *Artc1-*KO mice, increased immune responses and systemic inflammation with elevated serum levels of proinflammatory cytokines, e.g., TNF-*α*, decreased cardiac contractility and increased infarction induced by myocardial ischemia reperfusion primarily in male mice with upregulation of necroptosis marker protein RIP3, and reduced survival due to inflammation were seen compared to WT mice. Based on these findings, anti-ART1 agents can be therapeutic targets in cancer but beneficial effects may be limited by adverse effects such as inflammation in multiple organs and reduced cardiac contractility.

## Results

### Generation of Artc1-KO mice

*Artc1* knockout mice were generated using the CRISPR/Cas9 system. Three different lines of *Artc1*-KO mice were generated with three slightly different deletions in Exon 2 and 3 of the *Artc1* gene (Supplemental Figure 1A). Genotypes were confirmed by PCR (Supplemental Figure 1B) and Sanger sequencing. As expected in *Artc1*-KO mice, anti-ARTC1 antibodies did not detect the ARTC1 protein in *Artc1*-KO mice by Western Blotting (Supplemental Figure 1C).

### Reduced baseline myocardial contractility was seen in Artc1-KO male mice

Under basal conditions, 8-month-old *Artc1*-KO and *Artc1*-heterozygous (HT) male but not female mice, showed reduced myocardial contractility, e.g., percent ejection fraction (%EF), by magnetic resonance imaging (MRI) (Figure 1A and Supplemental Movie 1) and echocardiography (Echo) in B-mode (Supplemental Movie 2) and M-mode (Figure 1B) compared to their wild-type counterparts. Left ventricular end-diastolic diameter (LVEDD) and end-systolic diameter (LVESD) of *Artc1*-KO and *Artc1*-HT male, but not female mice, was significantly greater than those in WT mice (Figure 1C). By echocardiography, %EF and percent fractional shortening (%FS) were significantly less in *Artc1*-KO and *Artc1*-HT male mice than in WT male mice (Figure 1C), consistent with systolic dysfunction of *Artc1*-KO male mice observed by MRI. Cardiac fibrosis, a possible cause of systolic heart failure seen in *Arh1*-KO mice, was not observed in *Artc1*-KO but did in *Artc1*-KO */Arh1*-KO -double knockout mice (14.3 %, 4/28 male mice; 0%, 0/25 female mice), suggesting that the *Arh1* gene modulated the myocardial fibrosis seen in primarily male *Arh1*-KO mice (17). However, *Artc1*-KO male but not female mice exhibited increased heart weight/body weight (Supplemental Figure 3B).

**Figure 1.**
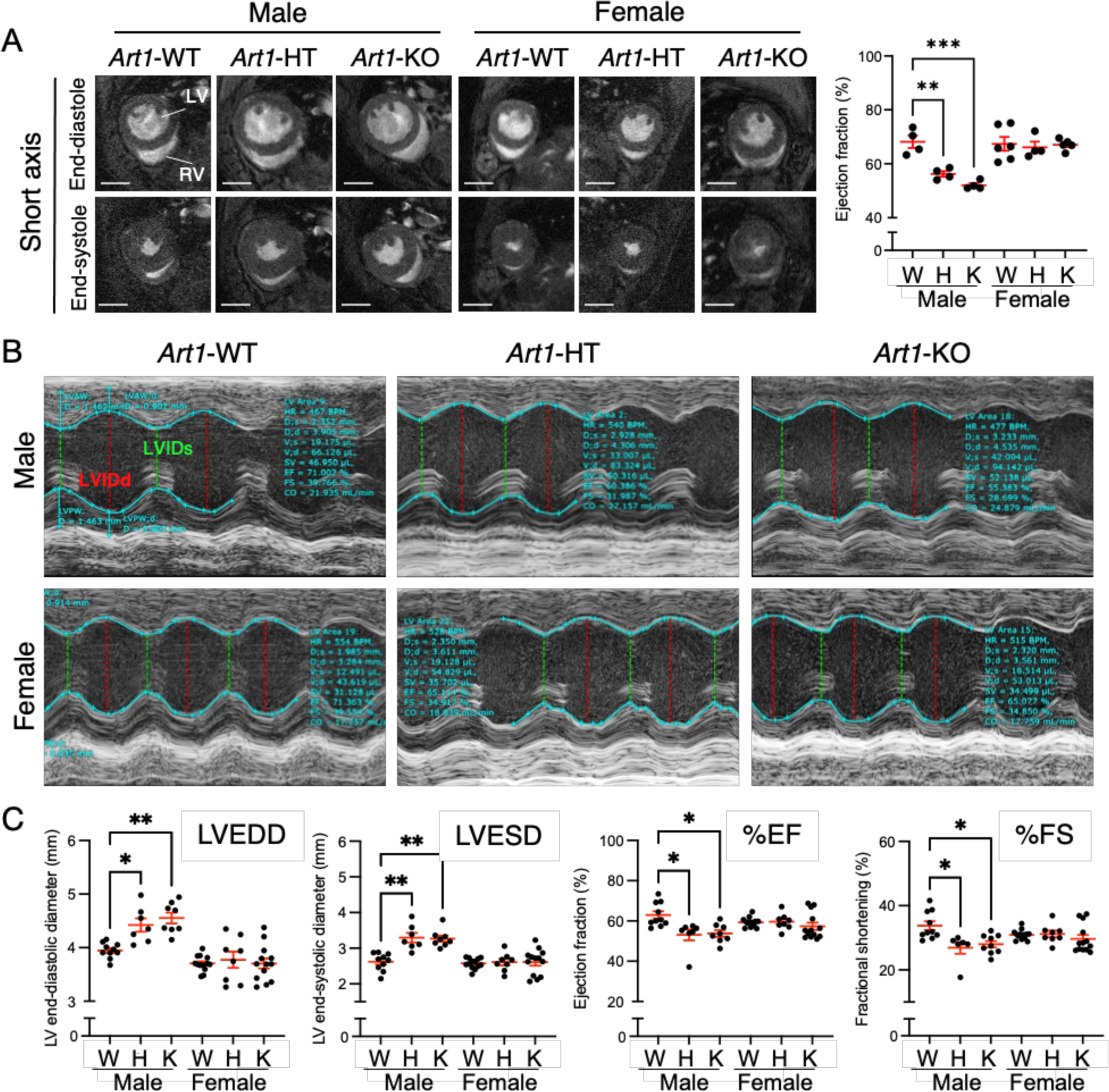
*Artc1*-KO and *Artc1*-heterozygous (HT) male mice showed systolic dysfunction. **(A)** *Artc1*-KO and *Artc1*-HT male showed decreased cardiac contractility by MRI. Cardiac contractility, e.g, percent ejection fraction (%EF), of *Artc1*-KO (male, *n*=4, 37.1 ± 1.4 weeks, %EF: 52.0 ± 1.6%; female, *n*=5, 39.9 ± 1.7 weeks, %EF: 67.1± 2.1%), *Artc1*-HT (male, *n*=4, 37.5 ± 0.6 weeks, %EF: 56.2 ± 2.0%; female, *n*=4, 38.8 ± 1.5 weeks, %EF: 66.1 ± 4.1%), and *Artc1*-WT (male, *n*=4, 39.0 ± 2.3 weeks, EF:68.2 ± 4.7%; female, *n*=6, 38.3 ± 1.8 weeks, %EF:67.4 ± 6.2%) mice were examined by MRI (Supplemental Movie 1). Typical images at end-diastole and end-systole by short axis are shown. Scale bar, 2 mm; Left ventricle, LV; right ventricle, RV. **(B)** Echocardiography showed decreased cardiac contractility in *Artc1*-KO and *Artc1*-HT males. Representative images by echocardiography in M-mode and movies of short axis (Supplemental Movie 2) are shown. (**C**) Left ventricle (LV) external end-diastolic (LVEDD) and end-systolic (LVESD) diameter were increased, and %EF and percent fractional shortening (%FS) were decreased in *Artc1*-KO (*n*=8, %EF: 53.8 ± 1.5%, %FS: 27.7 ± 1.0%) and *Artc1*-HT (*n*=7, %EF: 53.1 ± 2.3%, %FS: 26.9 ± 1.4%) male mice compared to *Artc1*-WT (*n*=10, %EF: 62.9 ± 1.9%, %FS: 33.8 ± 1.4%) male, but not female *Artc1*-KO (*n*=13, %EF: 57.4 ± 2.1%, %FS: 29.7 ± 1.4%) and *Artc1*-HT (*n*=8, %EF: 59.6 ± 1.3, %FS: 31.2 ± 0.9%) mice, compared to *Artc1*-WT (*n*=13, %EF: 59.3 ± 0.7%, %FS: 30.9 ± 0.5%) mice. Left ventricular internal dimension at systole, LVIDs; left ventricular internal dimension at diastole, LVIDd; *, *P* < 0.05; **, *P* < 0.01; ***, *P* < 0.1 vs. wild type of same gender by two-way ANOVA, followed by Bonferroni’s post hoc tests.

### Reduced cardiac contractility was seen in Artc1-KO mice with dobutamine stress test

Dobutamine (0, 10, and 40 μg/kg/min) was administered to evaluate cardiac function of *Artc1*-KO mice to mimic the effects of exercise on the heart. Myocardial contractility, e.g., percent ejection fraction (%EF), percent fractional shortening (%FS), showed a significant response to dobutamine in a dose-dependent manner, i.e., 10, and 40 μg/kg/min vs. 0 μg/kg/min dobutamine: respectively, WT male (*P* < 0.0001, *P* < 0.0001), *Artc1*-KO male (*P* = 0.002, *P* < 0.0001), WT female (*P* < 0.0001, *P* < 0.0001), and Artc1-KO female (*P* = 0.0024, *P* < 0.0001), regardless genotype or gender (Figure 2, A and B). *Artc1*-KO male mice consistently showed decreased %EF and FS at baseline and during low-dose and high-dose dobutamine infusion compared to *Artc1*-WT male mice. In female *Artc1*-KO mice, response to dobutamine-induced stress was reduced, i.e., %EF and %FS of *Artc1*-KO female mice were significantly less than those observed in wild-type female mice during low- and high-dose dobutamine infusion. These results are consistent with reduced cardiac function of *Artc1*-KO mice in both genders with dobutamine.

**Figure 2.**
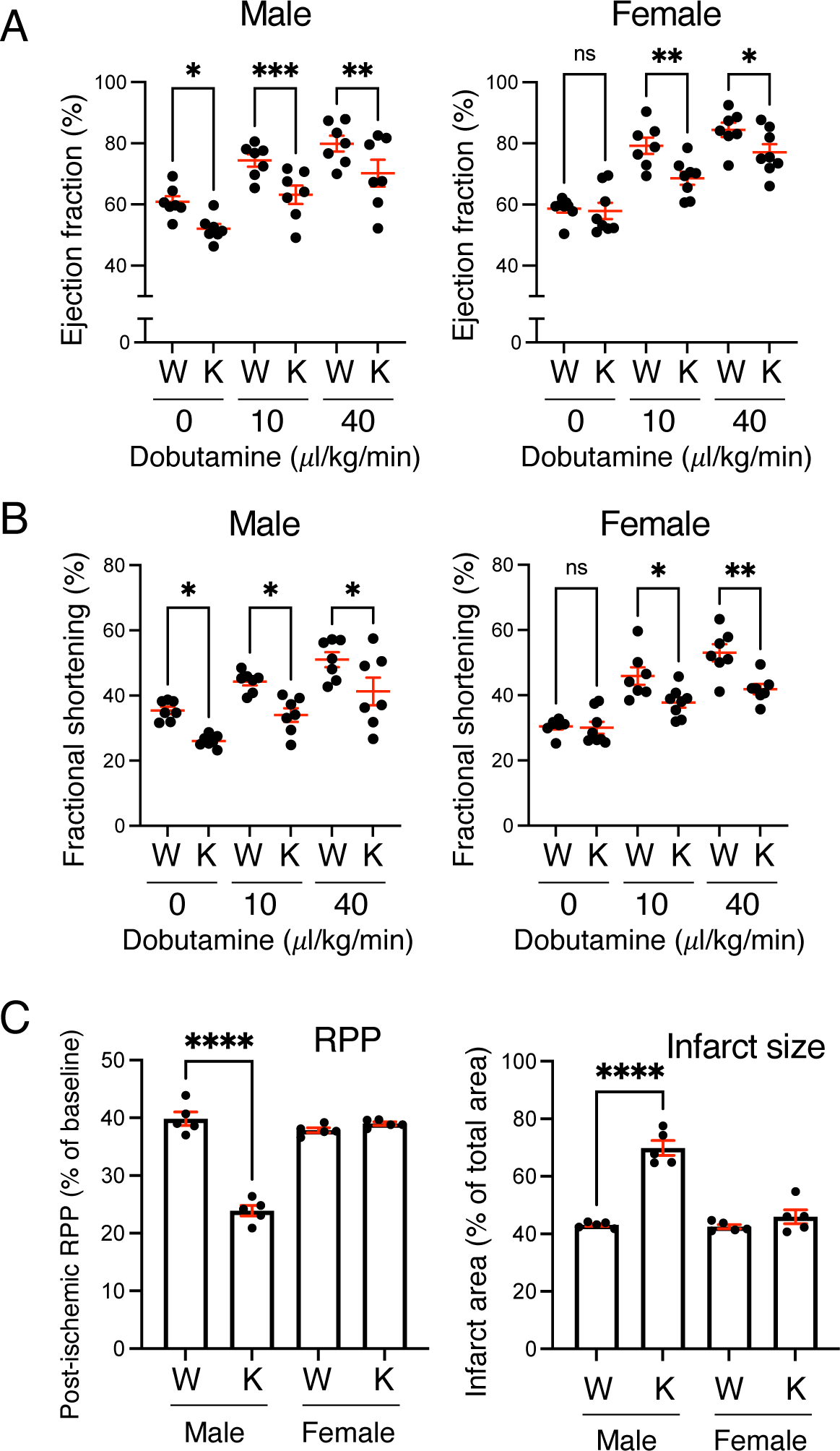
(A and B) Response to dobutamine-induced stress was reduced in *Artc1*-*KO* mice of both genders. During low- and high dose dobutamine (10 and 40 μl/kg/min) infusion, percent ejection fraction (%EF) and percent fractional shortening (% FS) were significantly lower in *Artc1*-KO mice (male, *n* = 7; female, *n* = 8) than in *Artc1*-WT (male, *n* = 7; female, *n* = 8) mice of same gender. **(C)** By Langendorff perfused heart model, *Artc1*-KO male, but not female mice, showed decreased RPP and increased infarct area. Isolated mouse hearts were perfused for 20 min prior to 25-min ischemia, followed by 90-min reperfusion. Recovery of post-ischemia left ventricular RPP and myocardial infarct size after ischemia/reperfusion in *Artc1*-KO mice (male, *n* = 5, 34.3 ± 2.5 weeks, 34.7 ± 1.7 g body weight (BW); female, *n* = 5, 35.7 ± 1.9 weeks, 25.0 ± 0.6 g BW) and in *Artc1*-WT littermates (male, *n* = 5, 35.3 ± 1.7 weeks, 34.2 ± 1.5 g BW); female, *n* = 5, 34.7 ± 1.5 weeks, 25.5 ± 0.7 g BW). Data are shown as mean ± SEM. *, *P* < 0.05; **, *P* < 0.01; ***, *P* < 0.001; ****, *P* < 0.0001 vs. wild type of same gender by two-way ANOVA, followed by Bonferroni’s post-hoc tests.

### In Langendorff perfused heart model, Artc1-KO male, but not female mice, showed increased infarct size and decreased rate pressure product (RPP)

Isolated mouse hearts were subjected to 20-min perfusion followed by 25-min ischemia/ 90-min reperfusion. There was no age (WT male, 35.3 ± 1.7 weeks; *Artc1*-KO male, 34.3 ± 2.5 weeks; WT female, 34.7 ± 1.5 weeks; *Artc1*-KO female, 35.7 ± 1.9 weeks) difference between genders and genotypes or body weight (WT male, 34.2 ± 1.5 g; *Artc1*-KO male, 34.7 ± 1.7 g; WT female, 25.2 ± 0.7 g; *Artc1*-KO female, 25.0 ± 0.6 g) between genotypes of same gender. Post-ischemic recovery of the RPP of *Artc1*-KO male mice (23.9 ± 0.8%) was decreased significantly compared to wild-type male mice (39.8 ± 1.9%) (Figure 2C). In female mice, no difference of post-ischemic RPP was observed between *Artc1*-KO and *Artc1*-WT mice, 39.0 ± 0.3% and 37.8 ± 0.4%, respectively. Infarct size, % of total area, after ischemia/reperfusion was greater in *Artc1*-KO male (69.8 ± 2.3%) than *Artc1*-WT male mice (43.1 ± 0.4%), but not between female mice (*Artc1*-KO female, 46.0 ± 2.2%; *Artc1*-WT female, 42.5 ± 0.7%) (Figure 2C).

### Tripartite motif 72 (TRIM72) was not ADP-ribosylated in Artc1-KO mouse heart lysates

Arginine-specific ADP-ribosylation of TRIM72 contributes membrane repair at the sites of injury caused by myocardial ischemia-reperfusion (17). ADP-ribosylated proteins, e.g., TRIM72, bound to Af1521, an ADP-ribose binding macrodomain, but not to an inactive mutant Af1521 macrodomain (38). As expected, wild-type Af1521 but not mutant Af1521 bound ADP-ribosylated TRIM72 in *Arh1*-KO mouse heart lysates (Supplemental Figure 2). Using this Af1521 pull-down system, ADP-ribosylated TRIM72 was not detected in *Artc1*-KO mouse heart lysates in vitro in both genders by immunoblot (Supplemental Figure 2), implying that *Artc1*-KO mice may have lost the functional ADP-ribosylation cycle involving TRIM72, thus resulting in myocardial injury (Figure 2C).

### ARTC1 contributed to cell migration during wound healing

To evaluate effect of ARTC1 on membrane repair and cell migration during wound healing, we generated *Artc1*-KO and *Artc1*-WT mouse embryonic fibroblasts (MEFs) (Figure 3A). *Artc1*-KO MEFs showed delayed scratch repair at 24 h compared to *Artc1*-WT MEFs (*P* < 0.0001) (Figure 3A).

**Figure 3.**
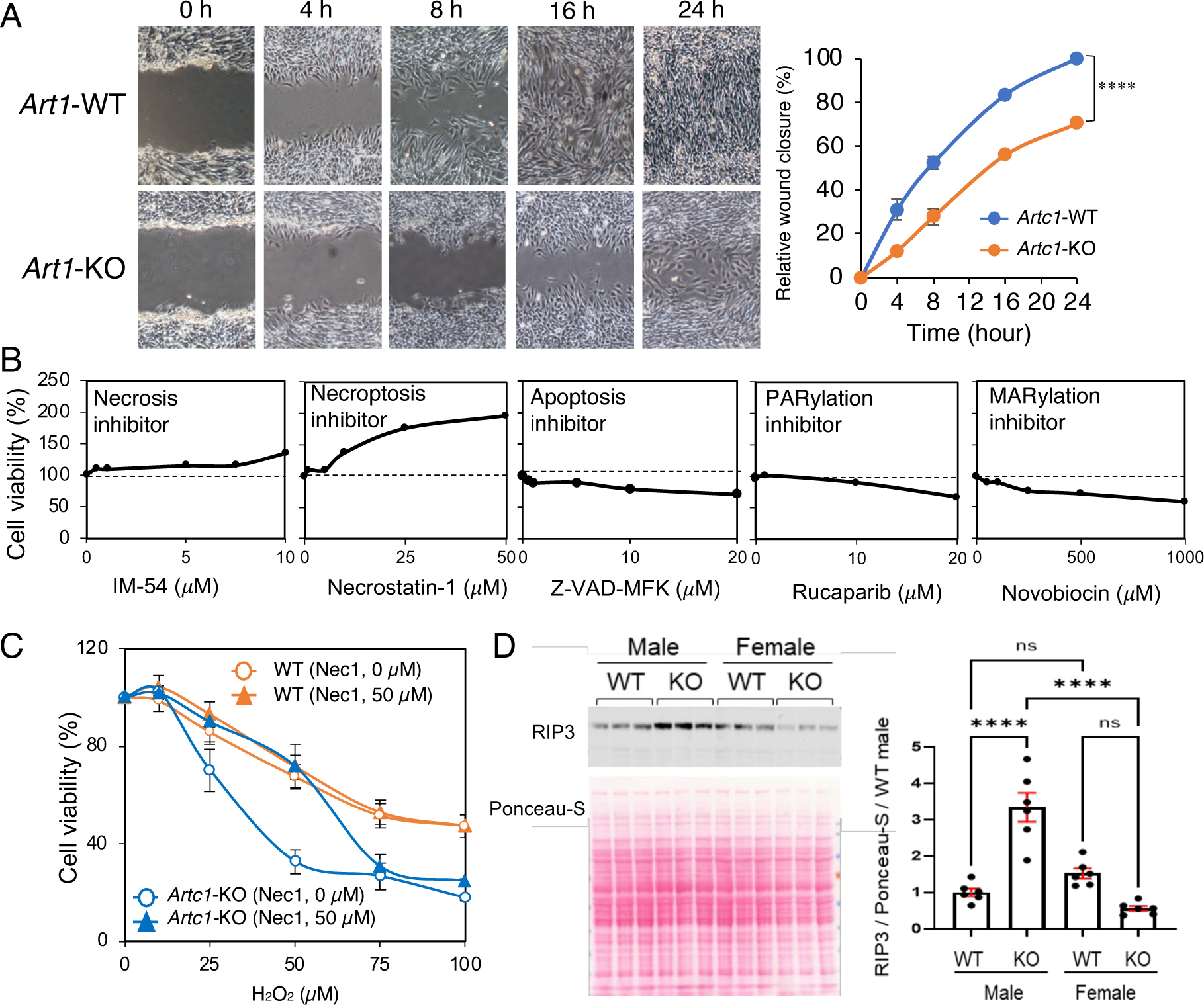
*Artc1* knockout upregulated necroptosis cell death pathway. **(A)** Scratch-wound-healing assays were performed with *Artc1*-KO and *Artc1*-WT MEFs. Images were taken at indicated time after wounding. Representative images are shown from three independent experiments. Wound area lacking cells was measured by ImageJ. Values are percentage of wound closure ± SEM per wound area at 0 h. *Artc1*-KO MEFs showed delayed wound closure compared to *Artc1*-WT MEFs. **(B)** Effects of necrosis (IM54), necroptosis (necrostain-1), apoptosis (Z-VAD-MFK), poly(ADP-ribosyl)ation (PARylation) (rucaparib), and mono-ADP-ribosylation (MARylation) (novobiocin) inhibitors on viability of *Artc1*-KO MEFs during exposure to 50 µM H_2_O_2_ -induced oxidative stress. *Artc1*-KO MEFs (1×10^4^ cells) were seeded on 96-well plates 24 h prior to 1-h treatment with inhibitors at indicated concentration, followed by oxidative stress with 50 µM H_2_O_2_. Cell viability of *Artc1*-KO MEFs was increased by 1.97 ± 0.08 times with 50 µM necrostain-1, but not necrosis and apoptosis inhibitors. Cell viability was measured by CCK-8 kit. **(C)** Necrostain-1 protected *Artc1*-KO MEFs from H_2_O_2_-induced oxidative stress. *Artc1*-KO MEFs were more sensitive to H_2_O_2_ dose-dependent oxidative stress compared to *Artc1*-WT MEFs. There were significant differences between *Artc1*-KO and *Artc1*-WT MEFs without necrostain-1 (*P* < 0.0001) and *Artc1*-KO MEFs with and without necrostain-1 (*P* = 0.007) by two-way ANOVA, followed by nonlinear regressions (GraphPad Prism). Data are average ± SEM of three independent experiments. (**D**) Hearts were harvested from mice without inflammation, e.g., ulcerative dermatitis. RIP3 protein levels were upregulated in *Artc1*-KO male hearts compared to *Artc1*-KO female and *Artc1*-WT male and female hearts. Representative Western blot and Ponceau-S images are shown from two individual experiments using three mice per mouse group. Statistical analysis was performed by two-way ANOVA, followed by Bonferroni’s post-hoc test by Prism 9.

### Necroptosis inhibitor increased cell viability of Artc1-KO mouse embryonic fibroblasts (MEFs) exposed to H_2_O_2_-induced oxidative stress

*Artc1*-*KO* MEFs showed decreased cell viability during H_2_O_2_-induced oxidative stress than did wild-type MEFs (*P* < 0.0001) (Figure 3C). Next, to define the responsible cell death pathway, MEFs were treated with inhibitors of necrosis (IM-54), necroptosis (necrostain-1), apoptosis (Z-VAD-FMK), poly-ADP-ribosylation (rucaparib), and mono-ADP-ribosylation (Novobiocin). Necrostatin-1 increased viability of *Artc1*-KO MEFs (*P* = 0.0007) exposed for 24 h to 50 µM H_2_O_2_-induced oxidative stress, whereas the other inhibitors did not improve viability (Figure 3B). Further, necrostatin-1 increased viability of *Artc1*-KO MEFs exposed to H_2_O_2_ at between 25 - 50 µM, *P* < 0.0001, for 24 h (Figure 3C). These results suggested that in MEFs ARTC1 participates in the necroptosis cell death pathway.

### RIP3 protein levels were higher in male Artc1-KO than in Artc1-WT mouse hearts

Therefore, next we compared RIP3 protein levels of *Artc1*-KO male and female mouse hearts. *Artc1*-KO male showed increased levels of RIP3 protein compared to *Artc1*-KO female and *Artc1*-WT mouse hearts (Figure 3D). Similar RIP3 levels were seen in WT female and male hearts (*P* = 0.85).

### Artc1 knockout and heterozygosity were associated with decreased tumor incidence

In contrast to the WT and *Arh1*-KO mice, male and female *Artc1*-KO did not develop tumors during a 24-month observation period, suggesting that the *Artc1* gene is a tumor promotor (Figure 4A). As we reported previously (16), *Arh1*-KO increased tumorigenesis (Figure 4C-b). Incidence of lymphoma and histiocytic sarcoma that was seen in wild-type mice (5.9%, 9/152; 5.9%, 9/152, respectively) was decreased in *Artc1*-heterozygous (3.0%, 9/303; 2.3%, 7/303, respectively) and homozygous (0%, 0/180; 0%, 0/180, respectively) *Artc1* knockout mice (Figure 4C-a). Increased incidence of tumorigenesis by *Arh1*-KO was dependent on a functional *Artc1* gene. Mice with decreased *Artc1* and *Arh1* expression, i.e., *Artc1*-KHT/*Arh1*-KO, *Artc1*-HT/*Arh1*-HT, were diagnosed only with lymphoma and histiocytic sarcoma (Figure 4C-c). Thus, regardless of *Arh1* genotype, *Artc1*-HT and *Artc1*-KO inhibited tumor development (Figure 4C-c), suggesting that ARTC1 is necessary to promote tumorigenesis.

**Figure 4.**
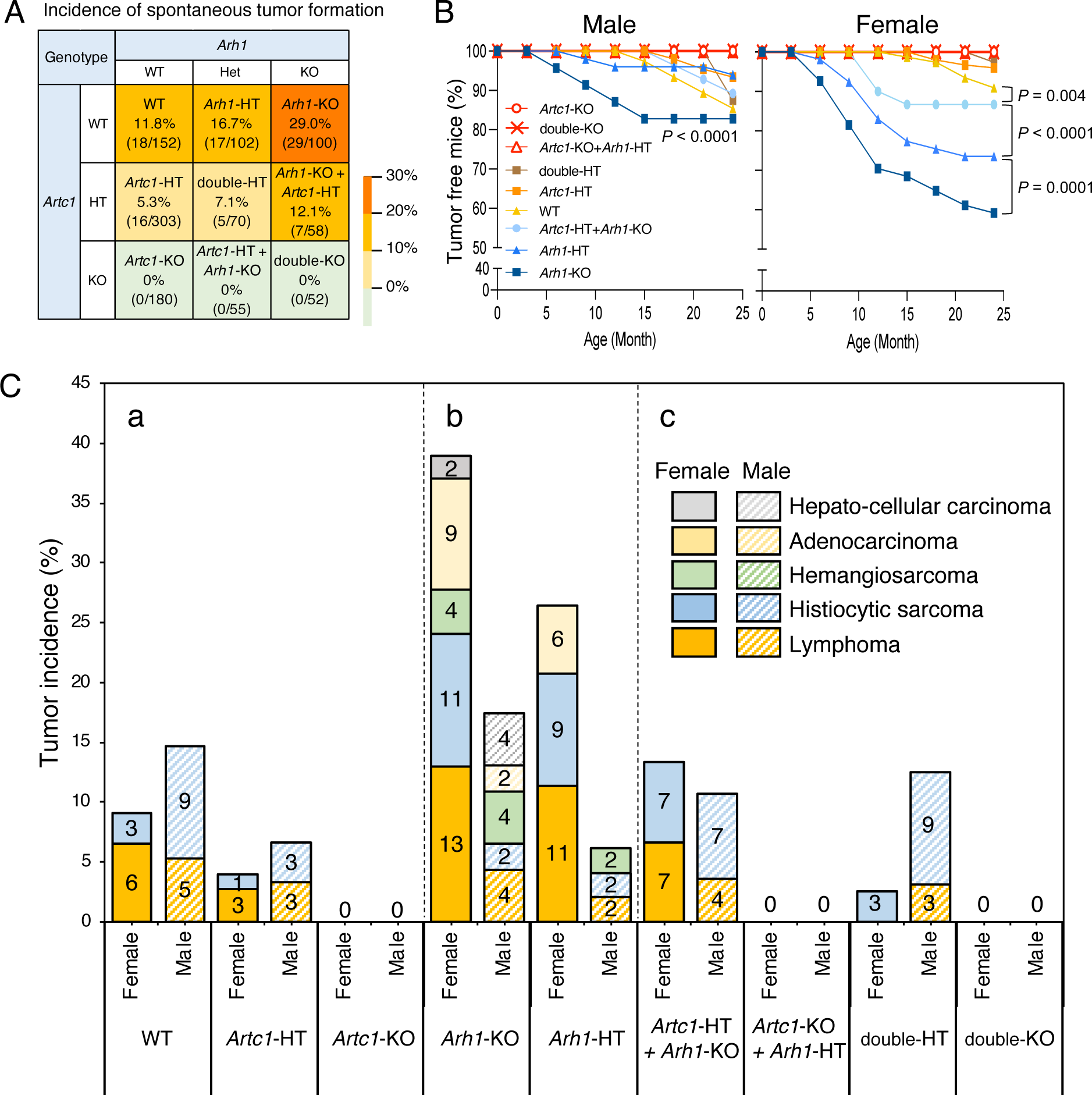
Effect of *Artc1* and *Arh1* genotypes on spontaneous tumor development. Spontaneous tumors of *Artc1* or *Arh1* homozygous knockout (KO), heterozygous (HT), *Artc1* plus *Arh1* homozygous knockout (double-KO), heterozygous (double-HT), and wild-type (WT) mice were observed for 24 months. **(A)** Incidence of spontaneous tumor in *Artc1* and *Arh1* mice, suggesting that *Artc1* and *Arh1* genes are tumor enhancer and suppressor, respectively. Number in parenthesis is number of mice bearing tumors/total mice. **(B)** Percentage of tumor-free mice by mouse age. Data for *Artc1*-KO, *Artc1*-KO+*Arh1*-HT, double-HT, and double-KO overlap each other. There were significant differences between genders in *Arh1*-KO (*P* < 0.0001) and *Arh1*-HT (*P* < 0.0001) mice, *Arh1*-KO male mice compared to WT mice of same gender as shown *P* value in Figure 4B, male. *Artc1*-deficient mice, e.g., *Artc1*-KO, *Artc1*-KO /*Arh1*-KO, *Artc1*-KO /*Arh1*-HT, did not develop tumors, were different from both male (*P* = 0.002) and female WT mice (*P* = 0.004). Statistical analysis was performed by two-way ANOVA followed by nonlinear regressions (GraphPad Prism). **(C)** Spontaneous tumors, e.g., lymphoma, histiocytic sarcoma, hemangiosarcoma, adenocarcinoma, hepato-cellular carcinoma, were evaluated by veterinary pathologist in a blinded manner. **(C-a)** *Artc1*-KO and *Artc1*-HT mice inhibited tumor formation. **(C-b)** Tumor incidence was increased in female mice by decreasing ARH1 level. **(C-c)** Depletion of *Artc1* suppressed *Arh1*-dependent tumor development that was enhanced by *Arh1* deficiency.

### Artc1 KO and heterozygosity delayed tumor initiation and progression induced by xenografts of Arh1-KO MEFs in nude mice

In mouse and cultured cell models, *Arh1-*KO and heterozygous (HT) mice and MEFs, respectively, developed tumors (16, 58). Subcutaneous injection of *Arh1*-KO, *Arh1*-HT, and *Arh1*-KO/*Art1*-HT MEFs in immune-deficient female nude mice lacking thymus gland by genetic mutation (athymic nude mice: *Foxn1^nucholer^* from Jackson Laboratory) resulted in tumor formation, whereas the other MEFs, e.g., *Artc1*-HT, *Artc1*-KO, *Arh1*-HT/*Artc1*-KO, *Arh1*-KO/*Artc1*-KO, were not tumorigenic (Figure 5A). Tumor doubling time of *Arh1*-KO /*Artc1*-HT MEFs was significantly (*P* < 0.0001) increased compared to *Arh1*-KO MEFs, implying that ARTC1 enhanced tumor progression by increasing cell proliferation (Figure 5B). Tumor formation by *Arh1*-KO/*Artc1*-HT MEFs was seen on average 10.5 days post-subcutaneous injection, which was slower than that of tumor formation by *Arh1*-KO MEFs where tumor formation was initiated in 24 h (Figure 5C). Thus, *Artc1*-KO and *Artc1*-HT delayed tumor initiation and growth of MEF xenografts in nude mice.

**Figure 5.**
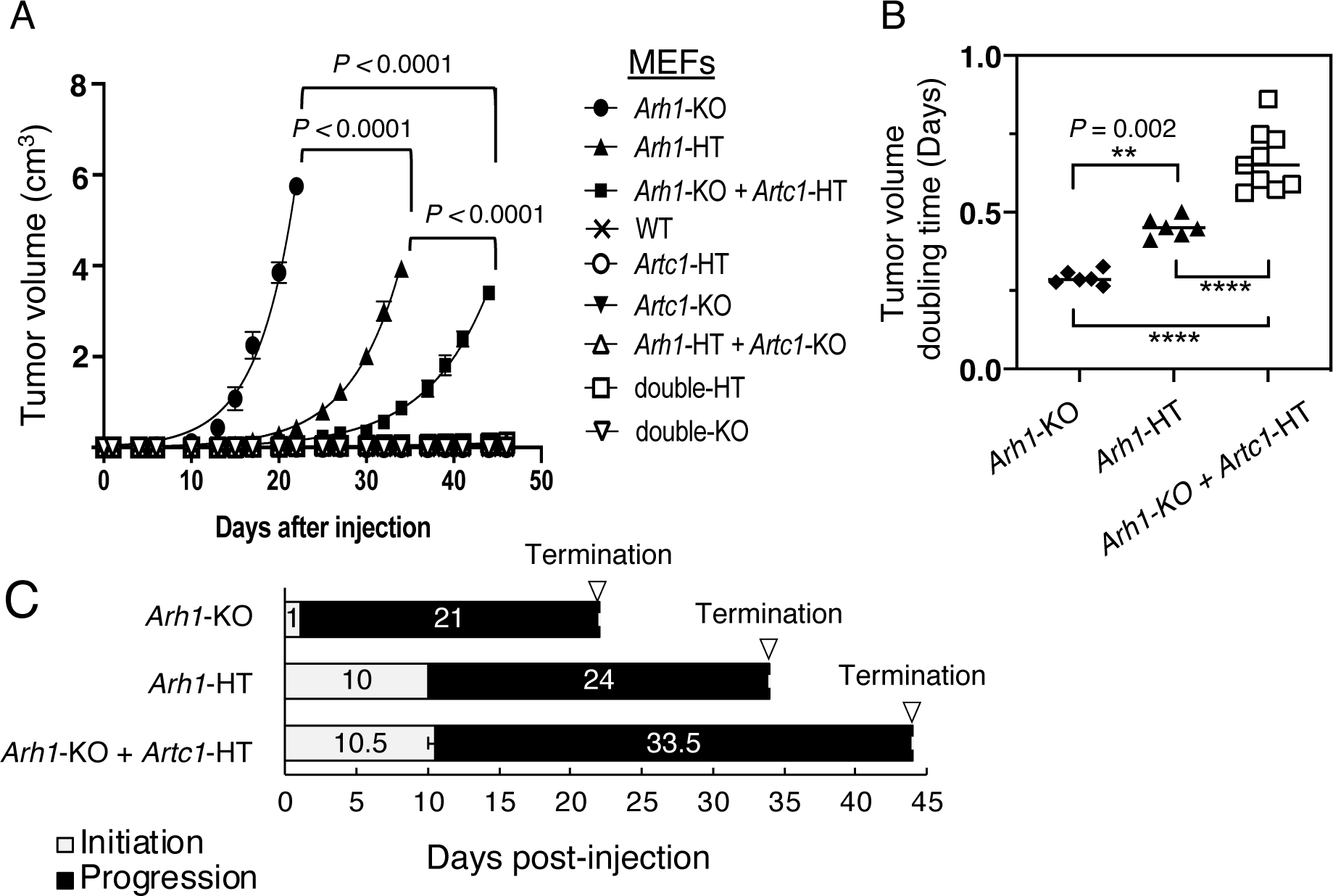
Effects of Arh1 and Artc1 genotypes on tumor growth in nude mice. MEFs were generated from following mice: Arh1 or Artc1 homozygous (KO), heterozygous (HT) knockout, Arh1+Artc1 homozygous-(double-KO) and heterozygous-(double-HT) knockout, Arh1-KO/Artc1-HT, Arh1-HT/Artc1-KO, wild-type (WT) mouse. MEFs were injected subcutaneously into nude mice. Tumor volume (cm^3^) was calculated by the modified ellipsoidal formula: Tumor volume = (length × width^2^) /2. (A) *Arh1*-KO, *Arh1*-HT, and *Arh1*-KO/*Artc1*-HT MEFs formed tumors in nude mice, but other MEFs did not. (B) Doubling times of tumor volume were significantly different among *Arh1*-KO, *Arh1*-HT, and *Arh1*-KO /*Artc1*-HT MEFs. (C) There were significant differences in tumor initiation time, i.e., the time when tumor appears, between *Arh1*-KO vs. *Arh1*-HT (*P* < 0.0001) or *Artc1*-HT/*Arh1*-KO (*P* < 0.0001), and in tumor progression time, i.e., the time from initiation time to the end point when tumor grew up to 2 cm, between *Arh1^−/−^* vs. *Arh1*-HT (*P* < 0.0001) or *Artc1*-HT*/Arh1*-KO (*P* < 0.0001) and *Arh1*-HT vs. *Artc1*-HT /*Arh1*-KO (*P* < 0.0001). For statistical analysis, two-way ANOVA followed by Bonferroni’s post-hoc test, or linear or nonlinear regression was carried out by GraphPad Prism. **, *P* < 0.01; ****, P < 0.0001.

### ARTC1 in the microenvironment increased tumor growth seen by subcutaneous injection of Arh1-KO MEFs

Compared to *Artc1*-WT mice, both male and female *Artc1*-HT and *Artc1*-KO mice showed decreased incidence of tumor formation following subcutaneous injection of *Arh1*-KO MEFs (Figure 6, A and B). Tumor incidence at day 37 following subcutaneous injection of *Arh1*-KO MEFs was 1.8 times greater (*P =* 0.026) in male *Artc1*-WT mice than in female *Artc1*-WT mice (Figure 6, A and B). Tumor growth rate did not differ among *Artc1* genotypes in male mice (Figure 6C), however, only in female *Artc1*-WT mice, did *Arh1*-KO MEFs develop tumors (Figure 6D). Overall, these results showed that the ARTC1 protein contributes to tumorigenicity by its presence in the tumor cell as well as in the microenvironment.

**Figure 6.**
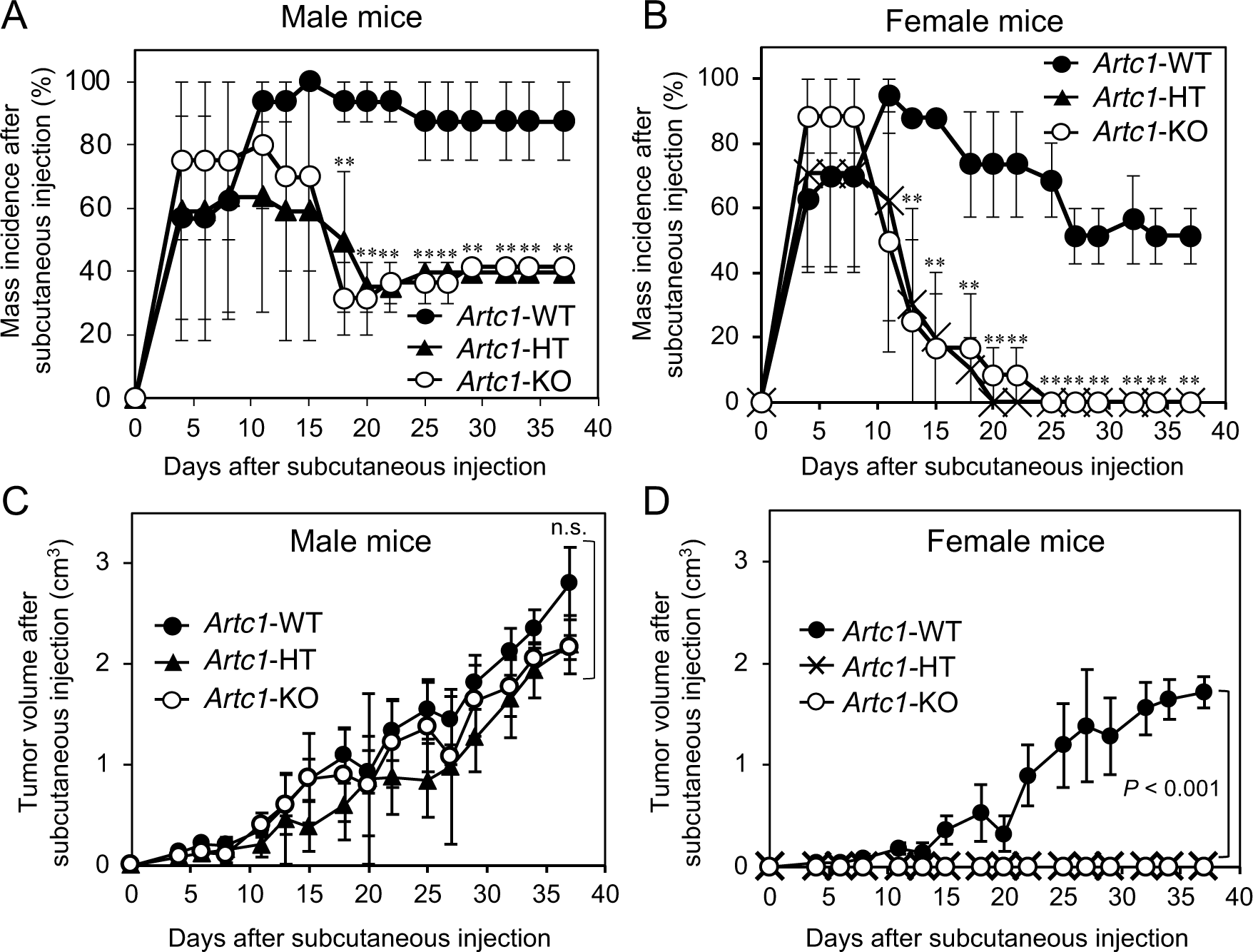
Artc1-*KO* and Artc1*-heterozygous (HT)* recipient mice decreased tumor formation induced by subcutaneous injection of Arh1-KO MEFs. Subcutaneous tumor growth was established by injecting tumorigenic *Arh1*-KO MEFs at the lower back of both male and female *Artc1*-WT, *Artc1*-HT, and *Artc1*-KO mice at 9- to 12-week-old. **(A** and **B)** Incidence of tumor was monitored for 37 days following subcutaneous injection of *Arh1*-KO MEFs into *Artc1*-WT, *Artc1*-HT, and *Artc1*-KO mice. In both genders, tumor incidence of *Arh1*-HT and *Arh1*-KO mice (male: after day 18, female: after day 13, respectively) was lower than seen with *Arh1*^+/+^ mice. Incidence rate of *Artc1* wild-type mice bearing tumors during a 37-day observation was 1.8 (*P* = 0.026) greater in male than female mice. *, significant difference versus wild-type mice of the same sex. **, *P* < 0.001. **(C** and **D)** Tumor volume was calculated by the modified ellipsoid formula (see Methods). Male mice bearing tumors showed similar tumor growth rate regardless genotypes (n.s., no significance), whereas in female mice, only wild type developed tumor. Male wild-type mice showed greater rate of tumor development than female wild-type (*P* < 0.0001). Statistical analyses including two-way ANOVA followed by linear and nonlinear regression and Bonferroni’s post-hoc test was carried out by GraphPad Prism.

### Enhanced immune cell accumulation was seen in tumors found in Artc1-KO and Artc1-HT mice

Areas of necrotic cells were found in both the periphery and center of tumors at day 8 after injection of *Arh1*-KO MEFs. Necrotic cell death was seen more frequently in female than male mice in the following order: *Artc1*-KO > *Artc1*-HT > *Artc1*-WT mice (Figure 7A, H&E stains). Infiltration of the *Arh1*-KO tumors by CD8^+^ cytotoxic T cells was dependent on *Artc1* genotypes, i.e., *Artc1*-KO and *Artc1*-HT mice showed enhanced infiltration of CD8^+^ lymphocytes in *Arh1*-KO MEFs tumors compared to *Artc1*-WT mice of same gender (Figure 7A and B). CD68^+^ M1 macrophages and CD209/DC-SIGN^+^ M2 macrophages were detected respectively in peripheral areas and necrotic areas of *Arh1*-KO MEFs tumors found in *Artc1*-KO and *Artc1*-HT mice (Figure 7A, CD209 and CD68). Tumors of *Artc1*-WT mice, however, exhibited less cell death as well as fewer macrophages than was seen in *Artc1*-HT and *Artc1*-KO mice (Figure 7A, CD209 and CD68). Levels of RIP3 protein, which is a promotor of necroptosis (60), a form of programmed necrosis, was increased in *Artc1*-KO tumors compared to *Artc1*-WT mice, suggesting that necroptosis was increased in tumors derived from *Arh1*-KO MEFs in *Artc1*-KO and *Artc1*-HT mice (Figure 7, C and D, RIP3). In contrast, levels of cleaved caspase-3 and cleaved PARP1, markers of apoptosis, were decreased in tumors formed in *Artc1*-KO mice (Figure 7, E and F). In agreement, apoptotic cells and cells in mitotic phase were more seen in wild-type and male mice compared to *Artc1*-KO mice (Figure 7, H&E stains). Tumors in nude mice derived from *Arh1*-KO MEFs did not have necrotic cells and infiltration by lymphocytes and macrophages that may cause necrotic cell death, at least until day11 (Supplemental Figure 3A). Taken together, these results suggest that absence of ARTC1 protein in *Artc1*-KO mice promoted infiltration of tumors with CD8^+^ lymphocytes and both M1 and M2 macrophages, leading to necroptosis of tumor cells.

**Figure 7.**
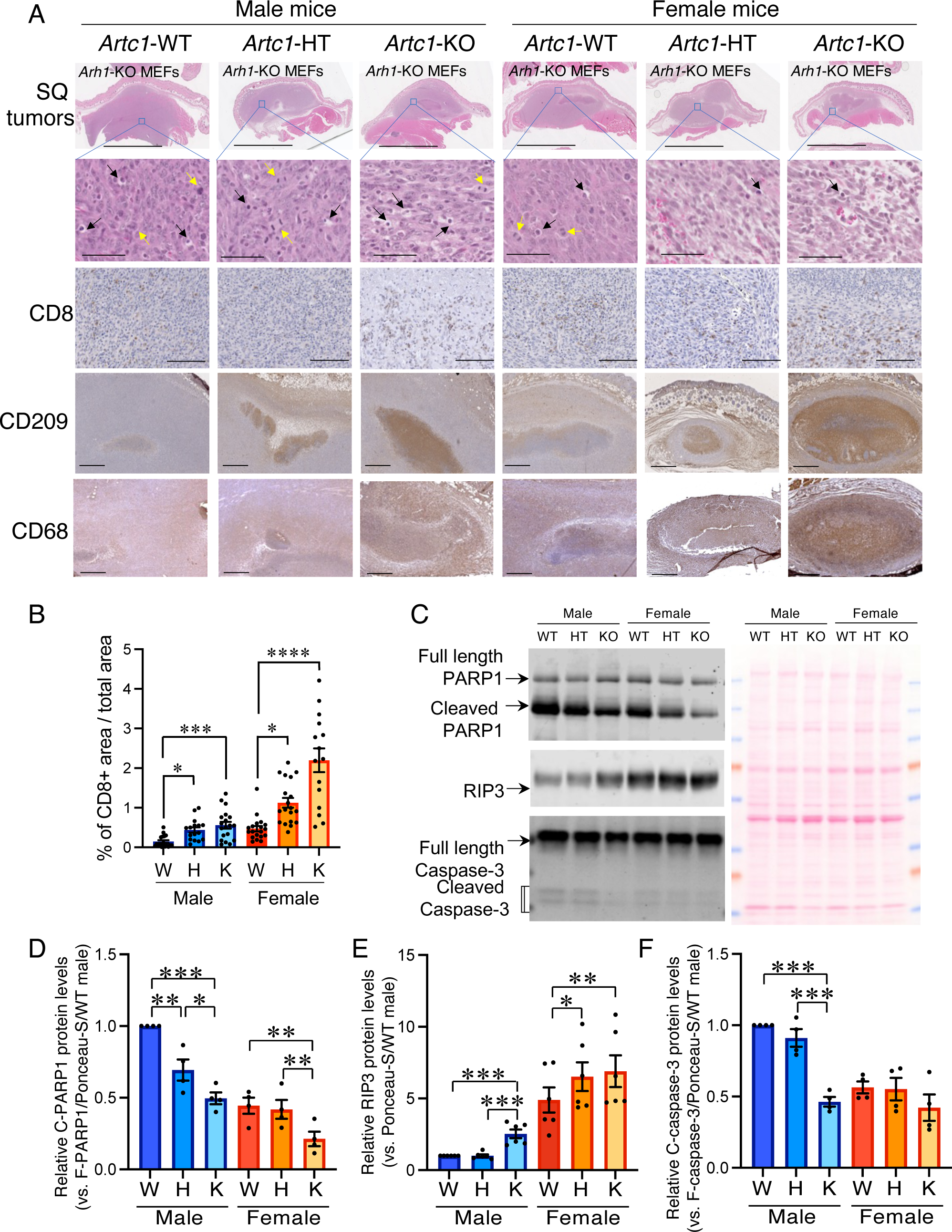
*Artc*1-dependent, tumor-infiltrating lymphocytes. **(A)** Necrotic cell death of tumors at day 8 formed by subcutaneous (SQ) injection of *Arh1*-KO MEFs in *Artc1*-WT, *Artc1*-HT, and *Artc1*-KO mice was evaluated with H&E staining of SQ tumors/ Typical images of H&E staining are shown (*n* = 5). Black and yellow arrows indicate, respectively, apoptotic cells and mitotic cells. Scale bar, 5 mm or 50 μm. Tumor-infiltrating lymphocytes and macrophages were stained with immunohistochemistry (IHC) against with anti-CD8 (cytotoxic T cell marker protein, 1:40 dilution), CD68 (M1 macrophage marker protein, 1:100 dilution), and CD209/DC-SIGN (M2 macrophage marker protein, 1:50 dilution) antibodies. **(B)** Percentage of CD8^+^ cells is presented as area of CD8^+^ cells / total area of tumor x 100. Quantification of IHC images was carried out by ImageJ. Typical images are shown (*n* = 4). Scale bar, 500 µm. **(C)** Apoptosis marker protein, e.g., PARP1, caspase-3, and necroptosis marker protein RIP3 levels of tumors at day 10 after subcutaneous injection of *Arh1*-KO MEFs in *Artc1*-WT, *Artc1*-HT, and *Artc1*-KO mice were detected by Western blotting with fluorescent-tagged secondary antibody. Ponceau-S stain was used as loading control of total protein. Typical images are shown from three individual experiments. **(D-F)** Quantification of protein levels compared to each genotype, *Artc1*-WT (W), *Artc1*-HT (H), *Artc1*-KO (K). Ratio of RIP3, cleaved PARP1 (C-PARP1)/full-length PARP1 (F-PARP1), and cleave caspase-3 (c-caspase-3)/caspase-3 protein levels were normalized by signal of Ponceau-S calculated by ImageJ, followed by protein levels of *Artc1*-WT male obtained by Odyssey CLx and Image Studio ver. 5.2. Statistical analysis was performed using two-way ANOVA followed by Bonferroni’s post-hoc test (GraphPad Prism 9). Significances are shown above bars: *, *P* < 0.05; **, *P* < 0.01; ***, *P* < 0.001.

### Artc1 knockout and heterozygosity was associated with increased immune response and inflammation

*Artc1*-KO and *Artc1*-heterozygous (HT) were associated with redness, swelling, pain, reduced food and water intake, which was evaluated by mouse facility staff who were blinded to mouse genotypes (61). Mice flagged for morbidity were evaluated for pathological assessment. Sick and found dead mice were sent for diagnostic necropsy. Immune response and inflammation were more frequent in *Artc1*-KO and *Artc1*-HT mice of both genders than in wild-type mice (Figure 8A) and were seen more in female than in male mice (Figure 8, B, C, and E). It was seen in an age-dependent manner in *Artc1*-KO, *Artc1*-HT, and *Artc1*-WT mice starting at 4, 11, and 16 months of age, respectively (Figure 8B). Of note, mice were separated in cages based on gender, not genotype. Thus, wild type, heterozygous, and homozygous knockout mice were kept in the same cages and exposed to the same conditions. Inflammation was observed in lung (e.g., edema, perivasculitis), kidney (e.g., nephritis), liver (e.g., hepatitis, necrosis), and skin (e.g., multifocal folliculitis, ulcerative dermatitis) (Figure 8, C and D), and increased immune response was seen in spleen (e.g., extramedullary) and lymph nodes (e.g., lymphadenopathy) (Figure 8, E and F). *Artc1/Arh1*-double KO female mice had reproductive system inflammation (16.7%). In rare instances (1.7%), bacterial thrombosis/endocarditis was observed in aortic valves of *Artc1*-KO hearts of both genders (Supplemental Figure 3C). CD8^+^ cytotoxic T cells, CD68^+^ M1 macrophages, and CD209^+^ M2 macrophages were increased in inflamed *Artc1*-KO spleen compared to wild-type spleen, whereas there was no difference of CD4^+^ helper T cell, CD19^+^ B cell, NKp46^+^ natural killer cells between *Artc1*-KO and *Artc1*-WT mouse spleen (Figure 8G). In agreement, plasma levels of pro-inflammatory cytokine TNF-*α*were increased in both male and female of *Artc1-*KO mice (Figure 9A). Due to increased immune response/inflammation, overall survival rate of male *Artc1*-KO and *Artc1*-HT mice showed reduced survival rate compared to WT mice (Figure 9B). *Artc1*-KO female mice was less than that of their heterozygous and WT counterparts (Figure 9C).

**Figure 8.**
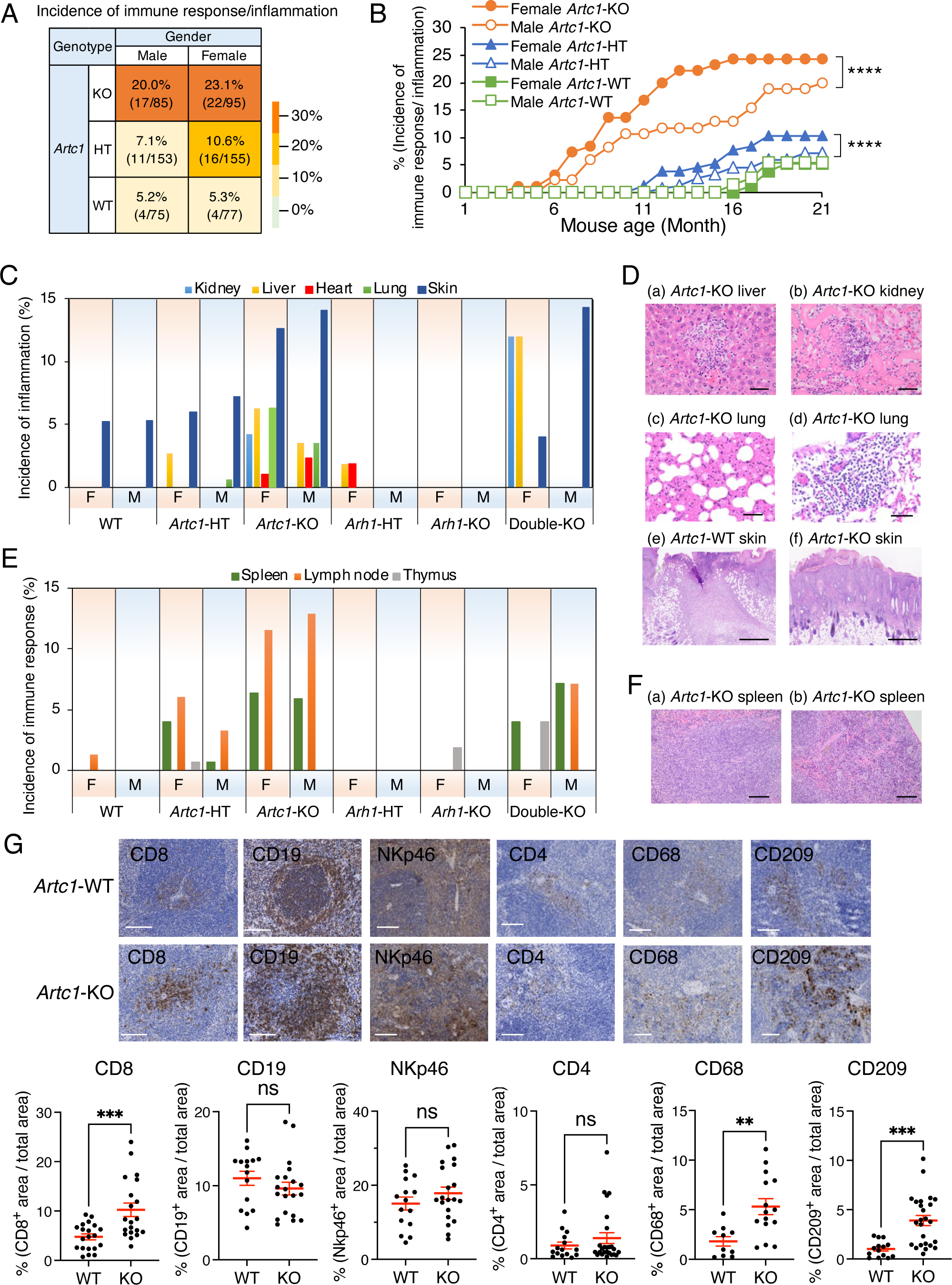
Immune response and inflammation in *Artc1* heterozygous (Het) and homozygous knockout (KO) mice. **(A)** Number of mice exhibiting immune response/inflammation. Number in parenthesis is number of mice with immune response/inflammation vs, total mice. Both male and female *Artc1*-KO mice showed higher incidence of immune response/inflammation than *Artc1*-WT (WT) mice. **(B)** Percentage of mice exhibiting immune response/inflammation by mouse age. There were significant differences between genders in *Artc1*-KO and *Artc1*-HT mice by nonlinear regressions (GraphPad Prism). ****, *P* < 0.0001. **(C)** Percentage of inflammation detected during a 21-month observation in multiple organs indicated above. *Artc1*gene mutations caused inflammation in multiple organs. M, male; F, female. **(D)** Representative H&E staining of *Artc1*-KO (a) liver with hepatitis (bar, 100 µm), (b) kidney with nephritis (bar, 100 µm), and lung with (c) edema (bar, 50 µm) and (d) perivasculitis (bar, 50 µm), and (e) WT and (f) *Artc1*-KO skin with ulcerative dermatitis (bar, 500 µm) exhibiting inflammation are shown. (E) Percentage of incidence of immune response detected during a 21-month observation in spleen, lymph node, and thymus. (F) Representative H&E staining of spleen (bar, 50 µm) exhibiting extramedullary in (a) white pulp and (b) red pulp. (**G)** CD8^+^ lymphocytes, and CD68^+^ and CD209^+^ macrophages accumulated more in spleen of *Artc1*-KO than *Artc1*-WT mice (*n* = 4 to 6). Immunohistochemistry-positive areas were detected by ImageJ and normalized by total area of each image. scale bar, 100 µm. Data are presented as mean ± SEM. ** *P* < 0.001, *** *P* < 0.0001, by two-tailed unpaired t test.

## Discussion

In this study, we identified *Artc1* genotype-dependent alterations in tumor formation, immune response/inflammation, cardiac contractility, and survival. We show that ARTC1 promotes tumor formation (Figure 4-7). ARTC1 protein was required in both tumor cells, i.e., MEFs (Figure 5), and in the tumor microenvironment, i.e., recipient mice, for tumor growth (Figure 6 and 7), suggesting that an anti-ARTC1 agent may have anti-cancer properties. In addition to suppressing tumor formation and growth, *Artc1*-KO was associated with decreased cardiac function (Figure 1 and 2), increased immune response/inflammation (Figure 8), and decreased survival (Figure 9, B and C). Of note, these potential adverse effects due to loss of ARTC1 may limit use of anti-ARTC1 agents as anti-cancer drugs.

**Figure 9.**
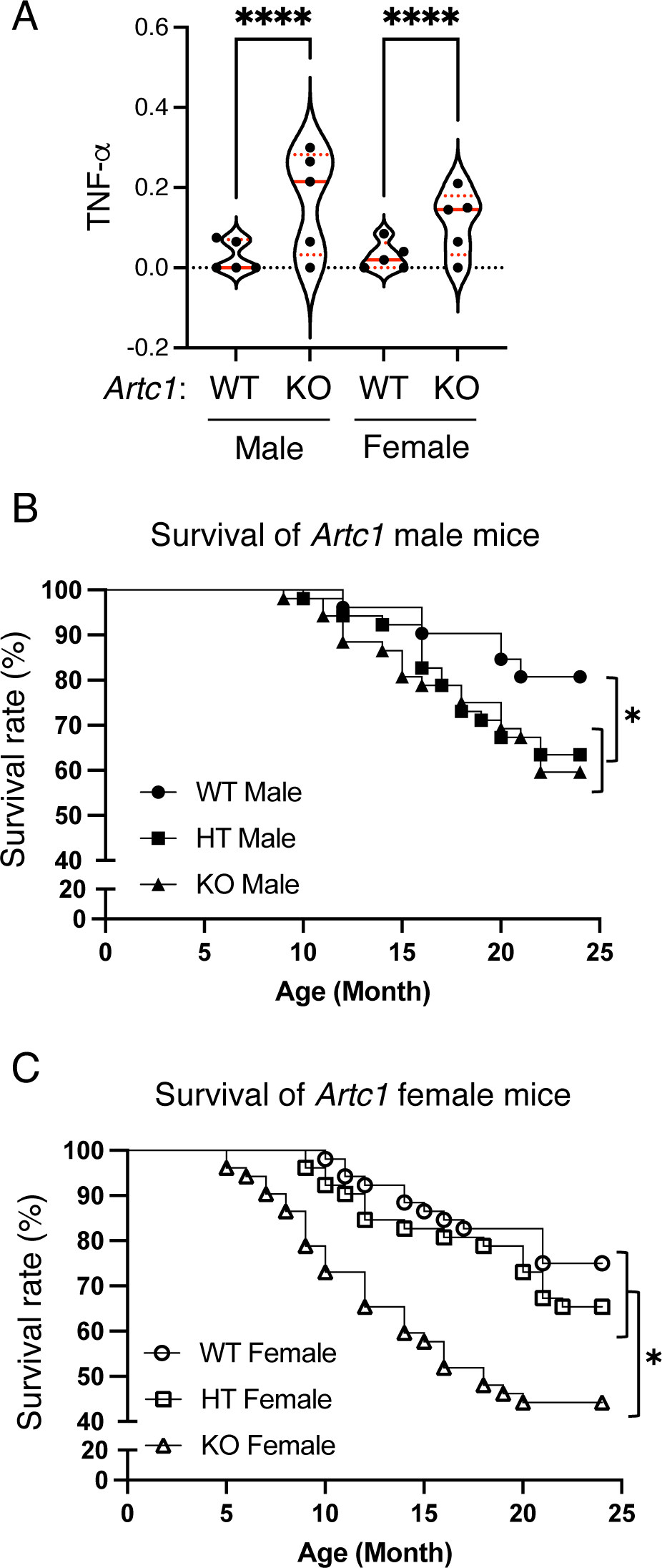
**(A)** IFN-*α* protein levels of *Artc1*-KO and WT mouse plasma. Mouse plasma with EDTA was collected from 5-month-old male and female of wild-type (*n* = 5), *Artc1*-KO (*n* = 5) mice. IFN-*α* protein levels were measured by mouse magnetic Luminex assay kit (R&D Systems) following the manufacturer’s instructions. Statistical analysis was performed by two-way ANOVA, followed by Bonferroni’s post-hoc test using GraphPad Prism. (**B** and **C**) Overall survival rate of *Artc1* mice. Mice were observed for 24 months (*n* = 52, each mouse group). Dead mice included those found dead (WT, 0%, HT, 0%, *Artc1*-KO male, 15.3%, *Artc1*-KO female, 7.4%) and euthanized based on 5-scale body condition assessment. Comparison of survival curves were performed by log-rank (Mante-Cox) test using Prism GraphPad. Significant differences (*, *P* < 0.05) were seen between genotypes, e.g., female KO vs WT, male KO vs WT, male Heterozygous (HT) vs WT, and gender, e.g., KO female vs KO male.

*Artc1* silencing inhibits proliferation, migration, and angiogenesis of HUVECs primary endothelial cells co-cultured with LoVo cancer cells transformed with *ARTC1*-shRNA (56). In agreement, overexpression of ARTC1 enhances tumorigenesis of CT26 colorectal carcinoma cells and promotes angiogenesis of HUVECs co-cultured with LoVo cells transfected with *ARTC1*-cDNA (56). In mouse xenograft model, a mouse small-cell lung cancer (SCLC) cell line, KP1, with shRNA targeting *Artc1*, showed increased apoptosis and infiltration with CD8^+^ T cells (13). These data suggest that tumor cells require ARTC1 for proliferation and migration. In this study, we used *Artc1*-HT and *Artc1*-KO recipient mice to examine whether the tumor microenvironment and the tumor cells required ARTC1 for tumor growth and metastasis. Based on the results of spontaneous tumor development by mice of different *Artc1* genotypes (Figure 4) as well as xenograft models with tumorigenic *Arh1*-KO MEFs (Figure 5-7), ARTC1 is required in both tumor cells, and in the tumor microenvironment, to promote tumorigenesis.

We reported previously that *Arh1*-KO and *Arh1*-HT mice developed spontaneous tumors more frequently in female than in male mice, suggesting that estrogen promotes tumorigenesis (62). In humans, however, males tend to develop cancer more frequently than females at any age, however, sex differences vary by cancer types (63, 64). Similar to human, *Artc1*-WT and *Artc1*-HT male mice exhibited a greater tumor incidence than seen in females (Figure 4C-a). *Artc1*-KO and *Artc1*-KO/*Arh1*-KO mice showed decreased tumor formation in both genders (Figure 4C), suggesting that *Artc1* regulates *Arh1* tumorigenicity and an anti-*Artc1* reagent may be effective in reducing tumors in both sexes in mice.

In contrast to the sex disparity in cancer, autoimmune disorders in humans are seen more frequently in females than in males (65). Females generally exhibit greater activity of innate and adaptive immune cells, e.g., monocytes, macrophages, than males in both humans and rodents, resulting in inflammatory diseases (66, 67). In this study, *Artc1*-KO female mice likewise exhibited greater frequency of immune response/inflammation in multiple tissues than did male mice (Figure 8), which resulted in lower overall survival rate in *Artc1*-KO female compared to *Artc1*-KO male mice (Figure 9, B and C).

In our previous report *Arh1*-KO male mice showed reduced cardiac contractility by echocardiography and MRI. Further, a subset (22%) of *Arh1*-KO male mice showed cardiac fibrosis. However, both males and females *Arh1*-KO mice showed increased myocardial infarction in an ex vivo Langendorff model (17). Therefore, we tested whether *Artc1*-KO mice both male and female were susceptible to ischemia-reperfusion induced myocardial injury in the Langendorff model. As expected, interrupting the cycle of arginine-specific mono-ADP-ribosylation by knocking out *Artc1* in males resulted in reduced cardiac contractility and increased myocardial infarction caused by ischemia/reperfusion-induced injury, while females showed normal cardiac

function and protection against ischemia/reperfusion-induced oxidative stress (Figure 1 and 2). Although plasma levels of pro-inflammatory cytokine TNF-*α*in both *Artc1*-KO male and female mice were upregulated compared to *Artc1*-WT mice of the same gender (Figure 9A), TNF-*α*-mediated necroptosis marker protein RIP3 levels were increased in *Artc1*-KO male mouse hearts but not in *Artc1*-KO female mouse hearts (Figure 3D). It has been suggested that estrogen has antioxidant properties (68), plays a cardioprotective role in women (69), and downregulates TNF-*α*-induced cytotoxicity in T cells (70). In agreement, RIP3 protein levels of *Artc1*-KO male hearts were greater than those of wild-type and *Artc1*-KO female hearts, suggesting that necroptosis cell death pathway via TNF-*α*-RIP3 was increased in *Artc1*-KO male hearts more than in *Artc1*-KO female hearts.

There are two confirmed in vivo substrates of ARTC1, e.g., human airway neutrophil peptide-1 (HNP-1) (9), and mouse heart tripartite motif-containing protein 72 (TRIM72) (17). ADP-ribosylated HNP-1 was isolated from bronchoalveolar lavage fluid (BALF) of patients with asthma and idiopathic pulmonary fibrosis. Di-ADP-ribosylation of HNP-1 on arginine residues by ARTC1 reduced both cytotoxicity of HNP-1 as well as its antibacterial activity, followed by non-enzymatic conversion of ADP-ribosylated arginines to ornithines leading to recovery of antibacterial activity, but not toxicity (15). In this study, bacterial infection was found in *Artc1*-KO mouse in skin e.g., ulcerative dermatitis (13.3%), and to a lesser extent in heart, e.g., endocarditis (1.7%), which was more frequent than in *Artc1*-WT mice, 1.3% and 0%, respectively (Figure 8C, Supplemental Figure 3C). Inflammation involved multiple organs, e.g., lung, kidney, liver, in both genders of *Artc1*-KO and *Artc1*-HT mice (Figure 8). ARTC1, however, is selectively expressed in immune cells, heart, and skeletal muscle, implying that circulating immune cells expressing ARTC1 may have an anti-inflammatory action in multiple organs. In addition, *Artc1*-KO mice had less granulation tissue in ulcer beds than wild-type mice (Figure 8D, skin). Formation of granulation tissue formation is important in healing of large ulcers by providing an antibacterial patch of extracellular matrix, e.g., collagen (71), blood vessels (72), and fibroblasts (73). Decreased granulation tissue formation of *Artc1*-KO mice may increase severity of bacterial infections in skin.

Anti-ARTC1 treatment with anti-ARTC1 antibody slowed tumor growth of KP1 cells without a side effect on organs and mouse body weight for up to three weeks (13). However, in our current study, *Artc1*-KO and *Artc1*-HT mice showed inhibition of tumor formation, increased systemic inflammation, decreased cardiac contractility, tumor infiltration by CD8^+^ T cells and macrophages and increased activation of a necroptotic cell death pathway. Necroptosis has morphological features of both necrosis and apoptosis, resulting in inflammation (74). Therefore, to reduce anti-ARTC1 reagent-induced inflammation, a necroptosis inhibitor may decrease the *Artc1*-KO-induced inflammatory response.

In conclusion, this study shows ARTC1 is required for tumor formation in both cells and their microenvironment, suggesting that anti-ARTC1 agents may be applicable as anti-cancer drugs. Furthermore, *Artc1-*KO male mice increased cardiac ischemia-reperfusion induced injury with upregulation of RIP3 protein levels, suggesting that anti-necroptosis agents may potentially be used for treating ischemia-reperfusion induced injury. Although, this study raises concerns on whether anti-ARTC1 agents may cause systemic inflammation and reduced cardiac contractility by TNF*α*-RIP3 induced necroptotic cell death pathway.

## Methods

### Animals

Experimental murine protocols (H-0127, H-0172, H-0177, and H-0271) were approved by the National Heart, Lung, and Blood Institute Animal Care and Use Committee. Mouse health was monitored by trained animal caretakers and facility veterinarians in a manner blinded to genotypes. Mouse pain assessment was performed using 5 scales: score of 0 - no pain or distress, score 1 - very minor pain or distress, score 2 - mild pain or distress, score 3 - moderate pain or distress that can meet criteria for euthanasia, score 4 - severe pain or distress that meets or exceeds criteria for euthanasia (61). Mice with scores higher than 2 or with loss of 20% body weight were euthanized, with histopathological evaluation of tumors and immune response/inflammation. Mice found dead were sent for diagnostic necropsies examination by board-certified veterinary pathologists. All mice were housed in facilities that were free from mouse parvovirus, but contained Helicobacter and murine norovirus. Heterozygous mice were bred to generate *Artc1*- and *Arh1*-KO, -heterozygous (HT), and -WT mice. Of importance regarding the effects of genotypes on immune response/inflammation, WT, HT, and KO mice of the same gender were kept in the same cage, hence, exposed to the same environment.

### Generation of CRISPR/Cas9 Artc1 knockout mice

The *Artc1*-KO mouse line was generated using CRISPR/Cas9 system (75) (Supplemental Figure 1A) . Briefly, two custom-designed, single guide RNAs (sgRNAs) were made by Thermo Fisher’s in vitro transcription service. One sgRNA, sgRNA-1 (Table 1), cuts shortly after the ATG initiation codon in Exon 2 (E2). The other sgRNA, sgRNA-2 (Table 1), cuts further downstream in Exon 3 (E3) (Supplemental Figure 1A). These sgRNAs (20 ng/*μ*l each) and Cas9 mRNA (50 ng/ *μ*l, purchased from Trilink Biotechnologies) were co-microinjected into the cytoplasm of zygotes collected from C57BL/6N mice (Charles River Laboratories). Injected embryos were cultured in M16 medium (MilliporeSigma) overnight in an incubator with 6% CO_2_ at 37 °C. The next morning, embryos that reached 2-cell stage of development were implanted into the oviducts of pseudopregnant surrogate mothers. Offspring born to the foster mothers were genotyped by PCR amplification of the targeted region using primers (Integrated DNA technologies, Coralville, Iowa) (Table 1, Supplemental Figure 1B). The PCR products were then sequenced using Sanger sequencing to select founder mice with frameshift deletions that caused premature translation termination. All three mutants, Mutant 1, 2, and 3, were used in a ratio of approximately 1:2:2 for 24-month observation (Figure 4, 8, and 9).

**Table 1.**
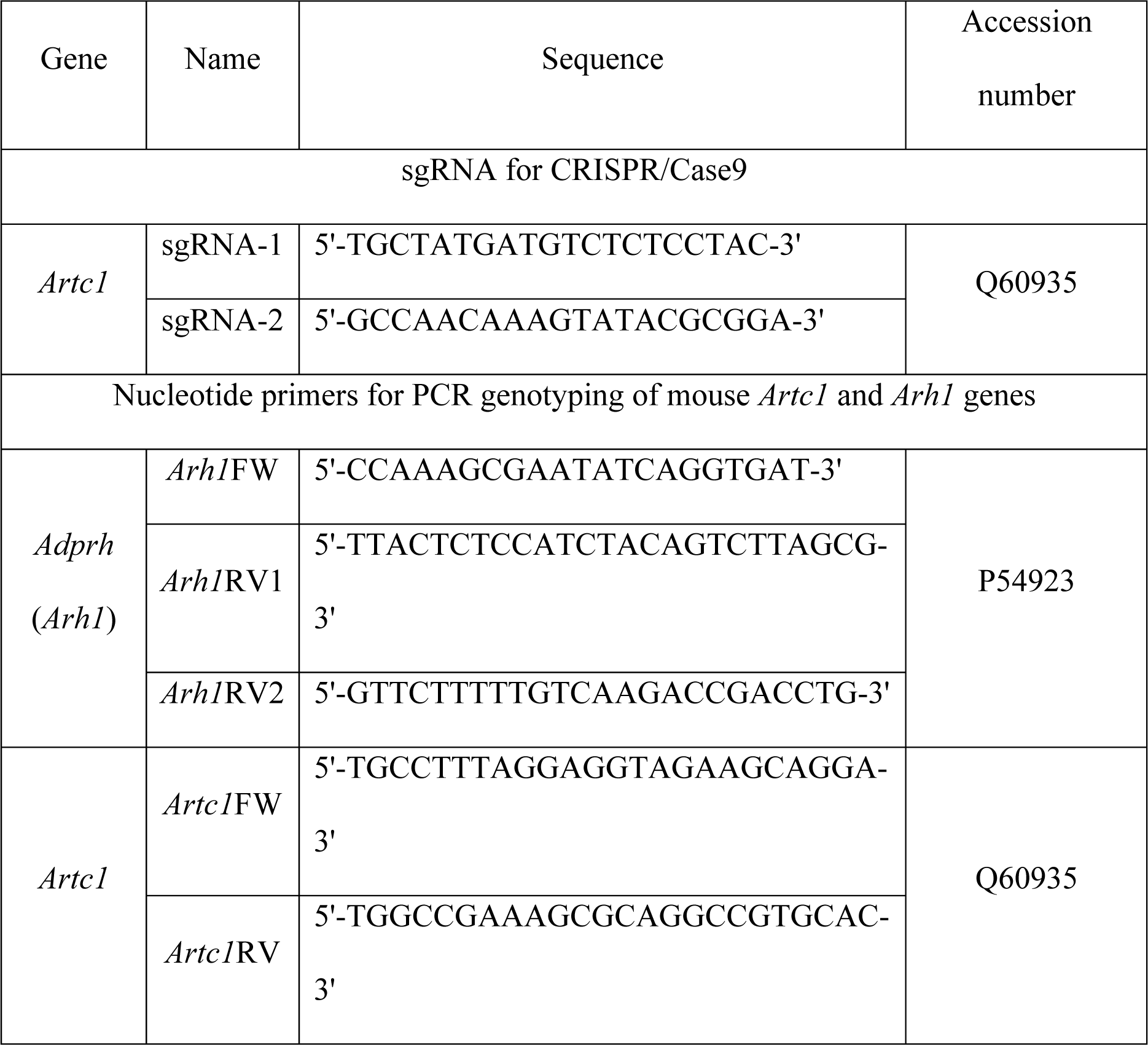
Sequence of single guide RNAs (sgRNAs) and PCR primers. Two sgRNAs, e.g., sgRNA-1, sgRNA-2, for mouse *Artc1* gene were used to generate *Artc1*-KO mice (Supplemental Figure 1A). Forward (FW) and reverse (RV) PCR primers of *Artc1* (Supplemental Figure 1B) and *Arh1* (14) were used for genotyping.

### Magnetic resonance imaging (MRI)

MRI was performed in a 7-Tesla MRI system with ParaVision software (Bruker, Billerica, MA). Mice were anesthetized with 2-3% isoflurane and placed in MRI coil, while monitoring ECG, respiratory rate, and body temperature (SA Instruments, Stony Brook, NY). Magnevist (Bayer Healthcare, Whippany, NJ) diluted 1:10 with sterile saline solution, was administered IP at 0.1 to 0.6 mmol/kg of mouse body weight. The sequence parameters were echo time (10 ms), size of field of view (30 x 30 mm or 48 x 28 mm), slice thickness (1 mm), and matrix (256 x 256 or 512 x 256).

### Baseline and dobutamine stress echocardiography

Mouse cardiac echocardiography was performed by an investigator blinded to mouse genotype. Mice were lightly anesthetized with isoflurane during examination and placed supine on a heated platform with a rectal temperature probe and electrocardiography (ECG) leads. Myocardial contractility was acquired in M-mode measurement of the left ventricle (LV) using the Vevo2100 ultrasound system (VisualSonics) with a 30-MHz ultrasound probe (VisualSonics, MS-400 transducer). After baseline scan, dobutamine (10, 40 µg/kg/min) was given to mice via the tail vein using an infusion syringe pump (Harvard Apparatus) at constant rate infusion (0.625 mg/ml dobutamine in saline containing 5% dextrose). Following the first infusion of dobutamine at low dose (10 µg/kg/min), when heart rate reached a steady state approximately 500 beats/min, infusion rate was increased to the high dose (40 µg/kg/min).

### Langendorff mouse heart perfusion model

Mice were anesthetized with sodium pentobarbital (50 mg/kg) by intraperitoneal injection and anti-coagulated with one-time heparin (50 μl, 1,000 U) administration into the inferior vena cava. Hearts were excised rapidly and placed in ice-cold Krebs Henseleit buffer (KH buffer) composed of 120 mM NaCl, 11 mM D-glucose, 25 mM NaHCO_3_, 1.75 mM CaCl_2_, 4.7 mM KCl, 1.2 mM MgSO_4_, and 1.2 mM KH_2_PO_4_ at pH 7.4. After cannulation of aorta with a Langendorff apparatus, the heart was perfused for 20 min in retrograde fashion with oxygenated (95% O_2_ / 5% CO_2_) KH buffer and maintained at 37°C at a constant pressure of 100 cm H_2_O. After 20-min perfusion, isolated hearts were subjected to no-flow ischemia for 25 min prior to 90 min of reperfusion. To monitor left ventricular contractile function, a latex balloon connected to a pressure transducer was inserted into the left ventricle of Langendorff-perfused hearts. Left ventricular developed pressure (LVDP) was recorded and digitized using a PowerLab system (ADInstruments). Recovery of post-ischemia left ventricular function: RPP (LVDP x heart rate) was used as a measure of function. Post-ischemic functional recovery was expressed as percentage of pre-ischemic RPP. For measurement of myocardial infarct size, after 90 min of reperfusion, the hearts were perfused with 1% 2,3,5-triphenyltetrazolium chloride (TTC; Millipore Sigma) dissolved in KH buffer and incubated in 1% TTC in KH buffer for 30 min at 37°C, followed by fixation in 10% formalin. Infarct size was expressed as the percentage of whole area of cross-sectional slices of mouse hearts.

### Cell lines

Mouse embryonic fibroblasts (MEFs) of all genotypes used in Figures 3 and 7-9, i.e., *Artc1*-KO, *Artc1*-HT, *Arh1*-KO, *Arh1*-HT, *Artc1*-KO/*Arh1*-KO, *Artc1*-HT/*Arh1*-HT, *Artc1*-KO/*Arh1*-HT, *Artc1*-HT/*Arh1*-KO, and wild-type, were produced by the same procedure described by Kato et al. (58). *Artc1*-HT and *Artc1*-KO MEFs were established using *Artc1* mouse strains of mutant 2 and 3 as indicated in Supplemental Figure 1.

### Cell viability assays

*Artc1*-KO and *Artc1*-WT MEFs were incubated on 96-well plates (1×10^4^ cells/well) for 24 h before exposure to H_2_O_2_ (0-100 µM, Sigma). Necrosis inhibitor IM-54 (0-10 µM, ChemCrus, EC_50_: 0.25 µM in HL-60 cells (76)), pan-caspase inhibitor Z-VAD-FMK (0-200 µM, Selleckchem, EC_100_: 50 µM in THP-1 cells (77)), necroptosis inhibitor necrostatin-1 (0-50 µM, MedChemExpress, EC_50_: 490 nM in 293T cells (74)), PARP inhibitor rucaparib (0-20 µM, MedChemExpress, EC50: 9 nM in HeyA8 cancer cells (78)), and mono-ADP-ribosylation inhibitor novobiocin (0-1 mM, Sigma, IC50: 700 µM in SkBr3 cells (79)) were added for 1 h prior to H_2_O_2_ exposure for 24 h. Cell viability was assessed using the CCK-8 kit (Dojindo, Rockville, MD) by measuring absorbance at 450 nm (SpectraMax, Molecular Devices, San Jose, CA) according to the manufacturer’s instructions.

### Antibodies

ARTC1 and TRIM72 antibodies were custom-made by 21st Century Biochemicals against mouse-specific peptides, CANSPLHKEFNAAVREA and CARLKTQLPQQKMQLQEA, respectively. Anti-PARP1, anti-cleaved PARP1, anti-RIP3, anti-caspase-3, anti-cleaved caspase-3, anti-CD8 (D4W2Z), and anti-CD19 (D4V4B) antibodies were from Cell Signaling. Anti-CD4 (RM4-5) and anti-NKp46 (PA5-102860) antibodies were from ThermoFisher. Anti-CD68 and anti-DC-SIGN/CD209 antibodies were from GeneTex. The secondary antibodies used for immunoblotting were anti-rabbit-IRDye 800CW antibodies and anti-mouse-IRDye 680RD antibodies from LI-COR.

### Subcutaneous injection

MEFs of all genotypes (1×10^6^ or 1×10^7^ cells per injection site as indicated in Figure legends) were injected subcutaneously at base of neck and hind limbs. After injection, the greatest longitudinal diameter (length) and transverse diameter (width) were measured using an electronic digital caliper to calculate tumor volume (cm^3^) by the modified ellipsoidal formula: Tumor volume = (length × width^2^) /2.

### Histopathology

Tissues were collected at the duty pathologist’s discretion for routine H&E histology. Complete gross examinations are performed on all submitted mice. All tissue samples were fixed in formalin and embedded in paraffin. Serial cross sections were stained with hematoxylin and eosin (H&E) for morphological evaluation. Immunohistochemistry (IHC) was performed with the avidin-biotin complex (ABC) staining method. Dilution of primary antibodies is indicated in the Figure legends.

### Immunoblot

Skeletal muscle lysates (30 µg of proteins) were resolved by SDS-PAGE (4-12% Bis-Tris NuPAGE gels). The proteins were transferred to nitrocellulose membranes, followed by Ponceau-S staining. Ponceau-S was removed by incubating with 0.05% Tween-20 in TBS (TBST) three times for 5 min. Membranes were blocked with 0.2% casein/TBST or 1% BSA/TBST for 1 h at room temp. After blocking, the membrane, and primary and fluorescent secondary antibodies, e.g., anti-rabbit-IRDye 800CW antibodies, anti-mouse-IRDye 680RD antibodies, were incubated with the same solution used for blocking the membranes. The proteins were detected by Odyssey CLX (LI-COR) and quantified by Image Studio software ver 5.2 (LI-COR).

### Cytokine quantification

Mouse plasma with EDTA was collected from male and female wild-type (20.7 ± 0.7 weeks, 22.2 ± 1.8 weeks, respectively), *Artc1*-KO (21.9 ± 2.1 weeks, 21.7 ± 1.5 weeks, respectively), *Arh1*-KO (22.9 ± 1.7 weeks, 21.9 ± 3.5 weeks, respectively), and *Artc1*-KO */Arh1*-KO (21.4 ± 0 weeks, 21.4 ± 0 weeks, respectively) mice. Mouse plasma cytokines were measured using mouse magnetic Luminex assay (LXSA, R&D Systems, MN, USA), following the manufacturer’s instructions. The assays included interferon (IFN)-*γ*, IFN-*α*, interleukin (IL)-1*α*, IL-1*β*, IL-2, IL-3, IL-4, IL-5, IL-6, IL-7, IL-10, IL-12 (p70), IL-13, IL-17A, chemokine CC ligand-2 (CCL2)/monocyte chemotactic protein (MCP)-1, CCL3/macrophage inflammatory proteins (MIP)-1*α*, CCL4/MIP-1*β*, CCL5/RANTES (regulated on activation normal T cell expressed), CCL11/Eotaxin, CXC motif chemokine ligand (CXCL)-1/GRO-*α*, CXCL2/GRO-*β*, CXCL13/ BLC (B-lymphocyte chemoattractant), epidermal growth factor (EGF), granulocyte colony-stimulating factor (G-CSF), granulocyte-macrophage colony-stimulating factor (GM-CSF), intercellular adhesion molecule 1 (ICAM-1)/CD54 (cluster of differentiation 54), matrix metalloproteinase-8 (MMP-8), tissue inhibitor of metalloproteinase-1 (TIMP-1), and hepatocyte growth factor (HGF). Cytokine levels were quantified in a Bio-Plex 200 System (Bio-Rad Laboratories) and analyzed by Bio-Plex Software (Bio-Rad Laboratories). Cytokine-concentration values below the lowest standard were extrapolated by the software and were included in the statistical analysis. Z-scores were obtained by converting the absolute values using the equation: Z-score = (sample concentration *−* mean of all samples) / standard deviation of all samples.

### Statistics

Data were presented as mean ± SEM. For analysis comparing two groups, two-tailed unpaired t test was performed using GraphPad Prism. Comparison among multiple data sets were first analyzed by two-way ANOVA, followed by linear, nonlinear regressions, or Bonferroni’s post-hoc test using GraphPad Prism. A p-value less than 0.05 was considered statistically significant.

### Study approval

All animal experiments were approved by the National Heart, Lung, and Blood Institute Animal Care and Use Committee: protocols H-0127, H-0172, H-0177, and H-0271.

## Supporting information

MRI videos of Artc1 mice

Echo videos of Artc1 mice

## List of Supplementary Materials

**Supplemental Movie 1.** *Artc1*-KO and *Artc1*-heterozygous (HT) male mice exhibited decreased myocardial contractility by MRI.

**Supplemental Movie 2.** *Artc1*-KO and *Artc1*-heterozygous (HT) male mice exhibited decreased myocardial contractility by echocardiography in B-mode.

**Supplemental Figure 1.** Generation of *Artc1*-KO mice.

**Supplemental Figure 2.** Pull-down assay of ADP-ribosylated TRIM72 in *Artc1*-KO heart lysates.

**Supplemental Figure 3.** A*r*h1-KO MEFs tumors formed in nude mice, mouse heart weight, and H&E stains of *Artc1*-KO heart with bacterial endocarditis.

## Acknowledgments

We thank Rodney L. Levine for valuable advice. We thank the staff at Diagnostic and Research Service Branch of Division of Veterinary Resources with necropsies and made slides. We appreciate the professional skills of Yukiko Kato (Pathology Core, National Heart, Lung, and Blood Institute, NIH) with histopathology-related experiments.

## Author contributions

J.K. and C.L. generated *Artc1*-knockout mice. J.K. observed effects of *Artc1* genotype on spontaneous incidence on tumor progression and inflammatory symptoms. D.A.S. performed echocardiography. M.J.L. performed MRI analysis. J.K. and H.O. performed subcutaneous injection of MEFs. J.S and E.M performed experiments involving the Langendorff model. Z.X.Y. assisted with processing of histopathological specimens and pathological evaluation. Z.X.Y. performed immunohistochemistry. P.D. and H.I.E. performed cytokine assays. H.I.E. performed cell viability assays, immunoblots, and statistical analysis. H.I.E. and JM helped design experiments, analyzed data and wrote the manuscript. All authors approved the manuscript.

---

**Supplemental Figure 1.**
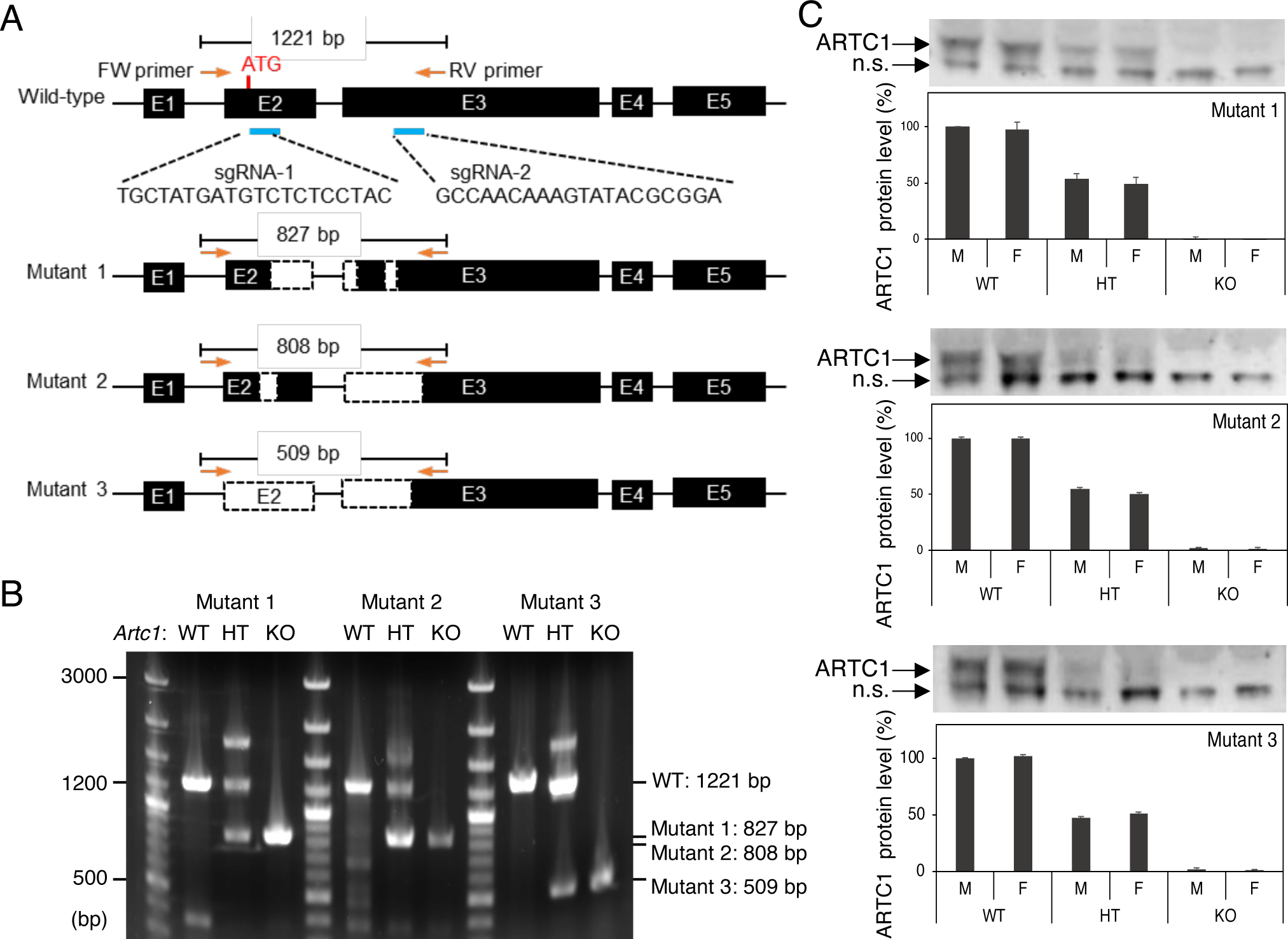
Generation of CRISPR/Cas9 *Artc1* knockout mice. **(A)** Two single-guide RNAs (sgRNAs) were designed, one cutting shortly after the translation initiation codon in Exon 2 (E2, the first coding exon) and the other one cutting in the coding region of Exon 3 (E3). These two sgRNAs and Cas9 mRNA were co-microinjected into zygotes collected from C57BL/6 mice, which were implanted into pseudopregnant foster mothers (CD-1 mice). C57BL/6J offspring were genotyped by PCR amplification and Sanger sequencing using PCR primers listed in Table 1. Three founder mice carrying different frame-shifting deletions were backcrossed to C57BL6J wild-type mice (Jackson Laboratory) for 8 generations. Heteroduplexes can form when there aren’t enough PCR reagents for amplifying all the denatured DNA molecules during the last a few cycles of the PCR reaction. In the presence of sufficient PCR reagents, all single-stand DNA will be converted into double-strand by Taq DNA polymerase during the extension step (72 °C), which results in the double-strand WT band and mutant band. However, when there aren’t enough PCR reagents (maybe one of the reagents, such as nucleotides or primer, is partially degraded), some of the denatured single-strand DNA cannot be converted to double-strand. After the temperature was lowered, the remaining single-strand DNA molecules anneal to each other, and some of the mutant DNA and WT DNA can anneal to each other forming heteroduplexes. **(B)** To maintain mouse colony, the genotypes of *Artc1* mutant mice were confirmed by PCR. **(C)** ARTC1 protein levels in the three mutant mouse lines, as analyzed by Western blotting. ARTC1 protein was undetectable in *Artc1*-KO mice in all three *Artc1* mutant lines, although a nonspecific protein (n.s.) was present in all mice. ARTC1 protein levels were reduced in heterozygotes for all three mutant lines.

**Supplemental Figure 2.**
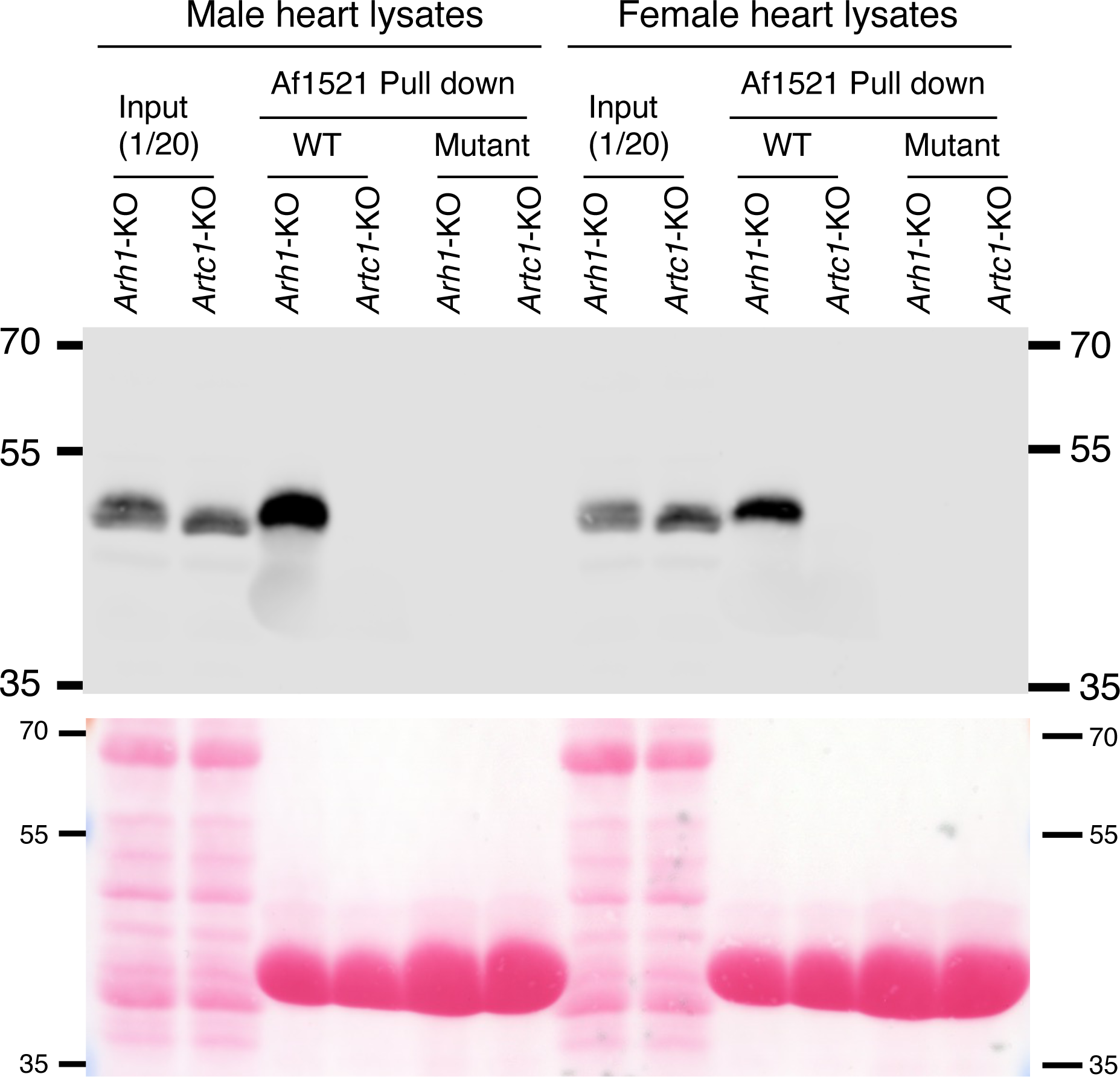
(**A**) ADP-ribosylation of membrane repair protein TRIM72/MG53 was not detected in *Artc1*-KO heart lysates from both male and female mice. *Artc1* -KO and *Arh1*-KO mouse hearts were lysed in 20 mM Tris-HCl / 20 mM NaCl / protease inhibitors, pH 7.5. The lysates were incubated with wild-type (WT) and inactive-mutant (Mutant) Af1521 macrodomain-GST bound to glutathione resin for 2 h at 4 °C, followed by 4-times washing with the binding buffer. The samples were applied to Western blotting with anti-TRIM72 antibody. TRIM72 protein was visualized with IRDye 800CW anti-rabbit IgG and detected by Odyssey imaging system. Ponceau-S staining was taken prior to blocking the membrane with 5% skim milk in TBST. Immunoblot and Ponceau-S images are representative of three experiments.

**Supplemental Figure 3.**
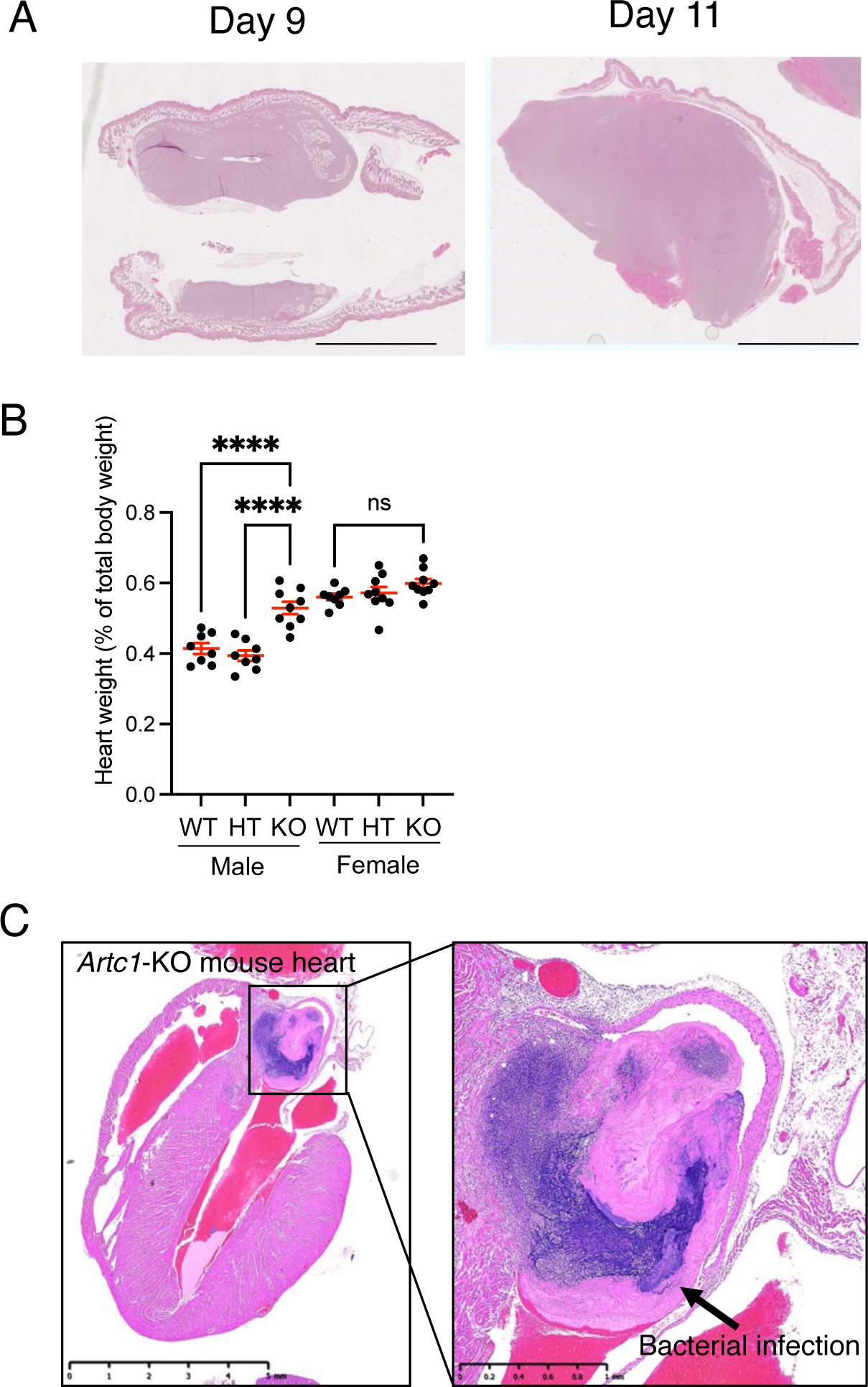
**(A)** Tumor induced by subcutaneous injection of *Arh1*-KO MEFs into nude mice. Typical images of H&E staining are shown (*n* = 4). Necrotic cells were not seen in tumors up to day 11 after subcutaneous injection of *Arh1*-KO MEFs. Scale bar, 5 mm. (**B**) percentage of heart-to-body weight of *Artc1*-WT (male, *n* =9, 29.6 ± 2.4 weeks; female, *n* = 10, 28.9 ± 1.7 weeks), *Artc1*-HT (male, *n* = 9, 30.3 ± 1.3 weeks; female, *n* = 10, 27.1 ± 1.9 weeks), and *Artc1-*KO (male, *n* = 11, 29.6 ± 2.4 weeks; female, *n* =10, 27.9 ±1.7 weeks) mice. Asterisk marks (****, *P* < 0.0001) are values significantly different by two-way ANOVA followed by Bonferroni’s post-hoc test. **(C)** Representative H&E staining of *Artc1*-KO mouse heart with bacterial infection. In rare cases (1.7%), both genders of *Artc1*-KO mice had a bacterial endocarditis in aortic valve.

## References

1. Fishman PH. Mechanism of Action of Cholera Toxin. ADP-ribosylating toxins and G proteins: Insights into Signal Transduction Moss and Vaughan, Eds 1990(American Society for Microbiology, Washington, D.C.):127–40.

2. Collier RJ. Effect of diphtheria toxin on protein synthesis: inactivation of one of the transfer factors. J Mol Biol. 1967;25(1):83–98.

3. Katada T, and Ui M. Direct modification of the membrane adenylate cyclase system by islet-activating protein due to ADP-ribosylation of a membrane protein. Proc Natl Acad Sci U S A. 1982;79(10):3129–33.

4. West RE, Jr., Moss J, Vaughan M, Liu T, and Liu TY. Pertussis toxin-catalyzed ADP-ribosylation of transducin. Cysteine 347 is the ADP-ribose acceptor site. J Biol Chem. 1985;260(27):14428–30.

5. Moss J, and Vaughan M. Mechanism of action of choleragen. Evidence for ADP-ribosyltransferase activity with arginine as an acceptor. J Biol Chem. 1977;252(7):2455–7.

6. Van Ness BG, Howard JB, and Bodley JW. ADP-ribosylation of elongation factor 2 by diphtheria toxin. NMR spectra and proposed structures of ribosyl-diphthamide and its hydrolysis products. J Biol Chem. 1980;255(22):10710–6.

7. Balducci E, Horiba K, Usuki J, Park M, Ferrans VJ, and Moss J. Selective expression of RT6 superfamily in human bronchial epithelial cells. Am J Respir Cell Mol Biol. 1999;21(3):337–46.

8. Donnelly LE, Rendell NB, Murray S, Allport JR, Lo G, Kefalas P, et al. Arginine-specific mono(ADP-ribosyl)transferase activity on the surface of human polymorphonuclear neutrophil leucocytes. Biochem J. 1996;315 (Pt 2):635–41.

9. Paone G, Stevens LA, Levine RL, Bourgeois C, Steagall WK, Gochuico BR, et al. ADP-ribosyltransferase-specific modification of human neutrophil peptide-1. J Biol Chem. 2006;281(25):17054–60.

10. Zolkiewska A, Nightingale MS, and Moss J. Molecular characterization of NAD:arginine ADP-ribosyltransferase from rabbit skeletal muscle. Proc Natl Acad Sci U S A. 1992;89(23):11352–6.

11. Liu ZX, Yu Y, and Dennert G. A cell surface ADP-ribosyltransferase modulates T cell receptor association and signaling. J Biol Chem. 1999;274(25):17399–401.

12. Fabrizio G, Di Paola S, Stilla A, Giannotta M, Ruggiero C, Menzel S, et al. ARTC1-mediated ADP-ribosylation of GRP78/BiP: a new player in endoplasmic-reticulum stress responses. Cell Mol Life Sci. 2015;72(6):1209–25.

13. Wennerberg E, Mukherjee S, Spada S, Hung C, Agrusa CJ, Chen C, et al. Expression of the mono-ADP-ribosyltransferase ART1 by tumor cells mediates immune resistance in non-small cell lung cancer. Sci Transl Med. 2022;14(636):eabe8195.

14. Kato J, Zhu J, Liu C, and Moss J. Enhanced sensitivity to cholera toxin in ADP-ribosylarginine hydrolase-deficient mice. Mol Cell Biol. 2007;27(15):5534–43.

15. Stevens LA, Barbieri JT, Piszczek G, Otuonye AN, Levine RL, Zheng G, et al. Nonenzymatic conversion of ADP-ribosylated arginines to ornithine alters the biological activities of human neutrophil peptide-1. J Immunol. 2014;193(12):6144–51.

16. Kato J, Vekhter D, Heath J, Zhu J, Barbieri JT, and Moss J. Mutations of the functional ARH1 allele in tumors from ARH1 heterozygous mice and cells affect ARH1 catalytic activity, cell proliferation and tumorigenesis. Oncogenesis. 2015;4:e151.

17. Ishiwata-Endo H, Kato J, Tonouchi A, Chung YW, Sun J, Stevens LA, et al. Role of a TRIM72 ADP-ribosylation cycle in myocardial injury and membrane repair. JCI Insight. 2018;3(22).

18. Okazaki IJ, and Moss J. Glycosylphosphatidylinositol-anchored and secretory isoforms of mono-ADP-ribosyltransferases. J Biol Chem. 1998;273(37):23617–20.

19. Ruf A, Mennissier de Murcia J, de Murcia G, and Schulz GE. Structure of the catalytic fragment of poly(AD-ribose) polymerase from chicken. Proc Natl Acad Sci U S A. 1996;93(15):7481–5.

20. Glowacki G, Braren R, Firner K, Nissen M, Kuhl M, Reche P, et al. The family of toxin-related ecto-ADP-ribosyltransferases in humans and the mouse. Protein Sci. 2002;11(7):1657–70.

21. Koch-Nolte F, Haag F, Braren R, Kuhl M, Hoovers J, Balasubramanian S, et al. Two novel human members of an emerging mammalian gene family related to mono-ADP-ribosylating bacterial toxins. Genomics. 1997;39(3):370–6.

22. Haag F, Koch-Nolte F, Kuhl M, Lorenzen S, and Thiele HG. Premature stop codons inactivate the RT6 genes of the human and chimpanzee species. J Mol Biol. 1994;243(3):537–46.

23. Weng B, Thompson WC, Kim HJ, Levine RL, and Moss J. Modification of the ADP-ribosyltransferase and NAD glycohydrolase activities of a mammalian transferase (ADP-ribosyltransferase 5) by auto-ADP-ribosylation. J Biol Chem. 1999;274(45):31797–803.

24. Leutert M, Menzel S, Braren R, Rissiek B, Hopp AK, Nowak K, et al. Proteomic Characterization of the Heart and Skeletal Muscle Reveals Widespread Arginine ADP-Ribosylation by the ARTC1 Ectoenzyme. Cell Rep. 2018;24(7):1916–29 e5.

25. Moss J, Jacobson MK, and Stanley SJ. Reversibility of arginine-specific mono(ADP-ribosyl)ation: identification in erythrocytes of an ADP-ribose-L-arginine cleavage enzyme. Proc Natl Acad Sci U S A. 1985;82(17):5603–7.

26. Hatakeyama K, Nemoto Y, Ueda K, and Hayaishi O. Purification and characterization of poly(ADP-ribose) glycohydrolase. Different modes of action on large and small poly(ADP-ribose). J Biol Chem. 1986;261(32):14902–11.

27. Okazaki IJ, and Moss J. Mono-ADP-ribosylation: a reversible posttranslational modification of proteins. Adv Pharmacol. 1996;35:247–80.

28. Munnur D, and Ahel I. Reversible mono-ADP-ribosylation of DNA breaks. FEBS J. 2017;284(23):4002–16.

29. Munnur D, Bartlett E, Mikolcevic P, Kirby IT, Rack JGM, Mikoc A, et al. Reversible ADP-ribosylation of RNA. Nucleic Acids Res. 2019;47(11):5658–69.

30. Fontana P, Bonfiglio JJ, Palazzo L, Bartlett E, Matic I, and Ahel I. Serine ADP-ribosylation reversal by the hydrolase ARH3. Elife. 2017;6.

31. Abplanalp J, Leutert M, Frugier E, Nowak K, Feurer R, Kato J, et al. Proteomic analyses identify ARH3 as a serine mono-ADP-ribosylhydrolase. Nat Commun. 2017;8(1):2055.

32. Hopp AK, and Hottiger MO. Uncovering the Invisible: Mono-ADP-ribosylation Moved into the Spotlight. Cells. 2021;10(3).

33. Rack JGM, Palazzo L, and Ahel I. (ADP-ribosyl)hydrolases: structure, function, and biology. Genes Dev. 2020;34(5-6):263–84.

34. Feijs KLH, Cooper CDO, and Zaja R. The Controversial Roles of ADP-Ribosyl Hydrolases MACROD1, MACROD2 and TARG1 in Carcinogenesis. Cancers (Basel). 2020;12(3).

35. Oka S, Kato J, and Moss J. Identification and characterization of a mammalian 39-kDa poly(ADP-ribose) glycohydrolase. J Biol Chem. 2006;281(2):705–13.

36. Moss J, Oppenheimer NJ, West RE, Jr., and Stanley SJ. Amino acid specific ADP-ribosylation: substrate specificity of an ADP-ribosylarginine hydrolase from turkey erythrocytes. Biochemistry. 1986;25(19):5408–14.

37. Ono T, Kasamatsu A, Oka S, and Moss J. The 39-kDa poly(ADP-ribose) glycohydrolase ARH3 hydrolyzes O-acetyl-ADP-ribose, a product of the Sir2 family of acetyl-histone deacetylases. Proc Natl Acad Sci U S A. 2006;103(45):16687–91.

38. Karras GI, Kustatscher G, Buhecha HR, Allen MD, Pugieux C, Sait F, et al. The macro domain is an ADP-ribose binding module. EMBO J. 2005;24(11):1911–20.

39. Martello R, Leutert M, Jungmichel S, Bilan V, Larsen SC, Young C, et al. Proteome-wide identification of the endogenous ADP-ribosylome of mammalian cells and tissue. Nat Commun. 2016;7:12917.

40. McClelland M, Sanderson KE, Spieth J, Clifton SW, Latreille P, Courtney L, et al. Complete genome sequence of Salmonella enterica serovar Typhimurium LT2. Nature. 2001;413(6858):852–6.

41. Allen MD, Buckle AM, Cordell SC, Lowe J, and Bycroft M. The crystal structure of AF1521 a protein from Archaeoglobus fulgidus with homology to the non-histone domain of macroH2A. J Mol Biol. 2003;330(3):503–11.

42. Rack JG, Perina D, and Ahel I. Macrodomains: Structure, Function, Evolution, and Catalytic Activities. Annu Rev Biochem. 2016;85:431–54.

43. Ame JC, Spenlehauer C, and de Murcia G. The PARP superfamily. Bioessays. 2004;26(8):882–93.

44. Pehrson JR, and Fried VA. MacroH2A, a core histone containing a large nonhistone region. Science. 1992;257(5075):1398–400.

45. Neuvonen M, and Ahola T. Differential activities of cellular and viral macro domain proteins in binding of ADP-ribose metabolites. J Mol Biol. 2009;385(1):212–25.

46. Slade D, Dunstan MS, Barkauskaite E, Weston R, Lafite P, Dixon N, et al. The structure and catalytic mechanism of a poly(ADP-ribose) glycohydrolase. Nature. 2011;477(7366):616–20.

47. Stevens LA, Kato J, Kasamatsu A, Oda H, Lee DY, and Moss J. The ARH and Macrodomain Families of alpha-ADP-ribose-acceptor Hydrolases Catalyze alpha-NAD(+) Hydrolysis. ACS Chem Biol. 2019.

48. Rosenthal F, Feijs KL, Frugier E, Bonalli M, Forst AH, Imhof R, et al. Macrodomain-containing proteins are new mono-ADP-ribosylhydrolases. Nat Struct Mol Biol. 2013;20(4):502–7.

49. Jankevicius G, Ariza A, Ahel M, and Ahel I. The Toxin-Antitoxin System DarTG Catalyzes Reversible ADP-Ribosylation of DNA. Mol Cell. 2016;64(6):1109–16.

50. Sharifi R, Morra R, Appel CD, Tallis M, Chioza B, Jankevicius G, et al. Deficiency of terminal ADP-ribose protein glycohydrolase TARG1/C6orf130 in neurodegenerative disease. EMBO J. 2013;32(9):1225–37.

51. Kleine H, Poreba E, Lesniewicz K, Hassa PO, Hottiger MO, Litchfield DW, et al. Substrate-assisted catalysis by PARP10 limits its activity to mono-ADP-ribosylation. Mol Cell. 2008;32(1):57–69.

52. Kasamatsu A, Nakao M, Smith BC, Comstock LR, Ono T, Kato J, et al. Hydrolysis of O-acetyl-ADP-ribose isomers by ADP-ribosylhydrolase 3. J Biol Chem. 2011;286(24):21110–7.

53. Stevens LA, Levine RL, Gochuico BR, and Moss J. ADP-ribosylation of human defensin HNP-1 results in the replacement of the modified arginine with the noncoded amino acid ornithine. Proc Natl Acad Sci U S A. 2009;106(47):19796–800.

54. Palavalli Parsons LH, Challa S, Gibson BA, Nandu T, Stokes MS, Huang D, et al. Identification of PARP-7 substrates reveals a role for MARylation in microtubule control in ovarian cancer cells. Elife. 2021;10.

55. Schleicher EM, Galvan AM, Imamura-Kawasawa Y, Moldovan GL, and Nicolae CM. PARP10 promotes cellular proliferation and tumorigenesis by alleviating replication stress. Nucleic Acids Res. 2018;46(17):8908–16.

56. Yang L, Xiao M, Li X, Tang Y, and Wang YL. Arginine ADP-ribosyltransferase 1 promotes angiogenesis in colorectal cancer via the PI3K/Akt pathway. Int J Mol Med. 2016;37(3):734–42.

57. Song GL, Jin CC, Zhao W, Tang Y, Wang YL, Li M, et al. Regulation of the RhoA/ROCK/AKT/beta-catenin pathway by arginine-specific ADP-ribosytransferases 1 promotes migration and epithelial-mesenchymal transition in colon carcinoma. Int J Oncol. 2016;49(2):646–56.

58. Kato J, Zhu J, Liu C, Stylianou M, Hoffmann V, Lizak MJ, et al. ADP-ribosylarginine hydrolase regulates cell proliferation and tumorigenesis. Cancer Res. 2011;71(15):5327–35.

59. Li Z, Yan X, Sun Y, and Yang X. Expression of ADP-ribosyltransferase 1 Is Associated with Poor Prognosis of Glioma Patients. Tohoku J Exp Med. 2016;239(4):269–78.

60. Declercq W, Vanden Berghe T, and Vandenabeele P. RIP kinases at the crossroads of cell death and survival. Cell. 2009;138(2):229–32.

61. Members ATF, Kohn DF, Martin TE, Foley PL, Morris TH, Swindle MM, et al. Public statement: guidelines for the assessment and management of pain in rodents and rabbits. J Am Assoc Lab Anim Sci. 2007;46(2):97–108.

62. Kato J, Bu X, and Moss J. Estrogen promotes tumorigenesis by ADP-ribosyl-acceptor hydrolase 1 (ARH1)-deficient cells and mice. Cancer Research. 2014;74(19).

63. Mitchell C, Richards S, Harrison CJ, and Eden T. Long-term follow-up of the United Kingdom medical research council protocols for childhood acute lymphoblastic leukaemia, 1980-2001. Leukemia. 2010;24(2):406–18.

64. Siegel RL, Miller KD, and Jemal A. Cancer Statistics, 2017. CA Cancer J Clin. 2017;67(1):7–30.

65. Libert C, Dejager L, and Pinheiro I. The X chromosome in immune functions: when a chromosome makes the difference. Nat Rev Immunol. 2010;10(8):594–604.

66. Melgert BN, Oriss TB, Qi Z, Dixon-McCarthy B, Geerlings M, Hylkema MN, et al. Macrophages: regulators of sex differences in asthma? Am J Respir Cell Mol Biol. 2010;42(5):595–603.

67. Boissier J, Chlichlia K, Digon Y, Ruppel A, and Mone H. Preliminary study on sex-related inflammatory reactions in mice infected with Schistosoma mansoni. Parasitol Res. 2003;91(2):144–50.

68. Manolagas SC. From estrogen-centric to aging and oxidative stress: a revised perspective of the pathogenesis of osteoporosis. Endocr Rev. 2010;31(3):266–300.

69. Ostadal B, and Ostadal P. Sex-based differences in cardiac ischaemic injury and protection: therapeutic implications. Br J Pharmacol. 2014;171(3):541–54.

70. Takao T, Kumagai C, Hisakawa N, Matsumoto R, and Hashimoto K. Effect of 17beta-estradiol on tumor necrosis factor-alpha-induced cytotoxicity in the human peripheral T lymphocytes. J Endocrinol. 2005;184(1):191–7.

71. Bailey AJ, Sims TJ, Le L, and bazin S. Collagen polymorphism in experimental granulation tissue. Biochem Biophys Res Commun. 1975;66(4):1160–5.

72. Weber K, and Braun-Falco O. Ultrastructure of blood vessels in human granulation tissue. Arch Dermatol Forsch. 1973;248(1):29–44.

73. Gabbiani G, Ryan GB, and Majne G. Presence of modified fibroblasts in granulation tissue and their possible role in wound contraction. Experientia. 1971;27(5):549–50.

74. Degterev A, Hitomi J, Germscheid M, Ch’en IL, Korkina O, Teng X, et al. Identification of RIP1 kinase as a specific cellular target of necrostatins. Nat Chem Biol. 2008;4(5):313–21.

75. Wang H, Yang H, Shivalila CS, Dawlaty MM, Cheng AW, Zhang F, et al. One-step generation of mice carrying mutations in multiple genes by CRISPR/Cas-mediated genome engineering. Cell. 2013;153(4):910–8.

76. Dodo K, Shimizu T, Sasamori J, Aihara K, Terayama N, Nakao S, et al. Indolylmaleimide Derivative IM-17 Shows Cardioprotective Effects in Ischemia-Reperfusion Injury. ACS Med Chem Lett. 2018;9(3):182–7.

77. Slee EA, Zhu H, Chow SC, MacFarlane M, Nicholson DW, and Cohen GM. Benzyloxycarbonyl-Val-Ala-Asp (OMe) fluoromethylketone (Z-VAD.FMK) inhibits apoptosis by blocking the processing of CPP32. Biochem J. 1996;315 (Pt 1)(Pt 1):21–4.

78. Hopkins TA, Ainsworth WB, Ellis PA, Donawho CK, DiGiammarino EL, Panchal SC, et al. PARP1 Trapping by PARP Inhibitors Drives Cytotoxicity in Both Cancer Cells and Healthy Bone Marrow. Mol Cancer Res. 2019;17(2):409–19.

79. Zhao H, Donnelly AC, Kusuma BR, Brandt GE, Brown D, Rajewski RA, et al. Engineering an antibiotic to fight cancer: optimization of the novobiocin scaffold to produce anti-proliferative agents. J Med Chem. 2011;54(11):3839–53.

